# Kidney toxicology of a novel compound Lithium Bis(trifluoromethanesulfonyl)imide (LiTFSI, ie. HQ-115) used in energy applications: an Epigenetic evaluation

**DOI:** 10.1101/2024.04.02.587863

**Authors:** Mia Sands, Xing Zhang, Joseph Irudayaraj

**Author notes:** These authors contributed equally to this work (Mia Sands, Xing Zhang).

## Abstract

Exposure to emerging energy-based environmental contaminants such as lithium bis(trifluoromethanesulfonyl)imide (LiTFSI), more commonly known as HQ-115, poses a significant threat to human health, yet its impact on kidney function and epigenetic regulation remains poorly understood. Here, we investigated the effects of LiTFSI exposure on kidney-related biochemical parameters, renal injuries, and epigenetic alterations in male CD-1 mice under both 14-day and 30-day exposure durations. Our study revealed that LiTFSI exposure led to changes in kidney-related biochemical indicators, notably affecting serum bicarbonate levels, while relative kidney weight remained unaffected. Histological analysis unveiled tubule dilation, inflammation, and loss of kidney structure in LiTFSI-exposed mice, alongside dysregulated expression of genes associated with inflammation, renal function, and uric acid metabolism. Epigenetic analysis further identified widespread DNA methylation changes in the two exposure regimes. Functional analysis revealed that differentially methylated regions are implicated in cell apoptosis and cancer-related pathways and are enriched with development-related transcription factor binding motifs, suggesting a potential mechanism of action that can lead to kidney injury. These findings underscore the intricate interplay between environmental exposures, epigenetic modulation, and kidney health, emphasizing the need for additional research to unravel precise mechanisms that can help in the development of targeted interventions to mitigate the adverse effects of LiTFSI exposure on human health.

**SYNOPSIS:** LiTFSI (HQ-115), an emerging environmental contaminant, impacts kidney health in male CD-1 mice by altering biochemical indicators, to result in renal injuries, and inducing epigenetic changes, highlighting environmental health concerns.

## 1. INTRODUCTION

Per-and polyfluoroalkyl substances (PFAS) encompass a class of synthetic organic compounds characterized by the substitution of hydrogen atoms in the alkyl structure with fluorine atoms. These characteristics result in PFAS possessing remarkable amphiphilic properties and robust carbon-fluorine (C-F) bonds, rendering them highly stable, hydrophobic, oleophobic, and characterized by low surface tension (Brusseau et al., 2020; Crone et al., 2019; Evich et al., 2022). Consequently, PFAS find widespread applications across various industries, including as lubricants, surfactants, refrigerants, adhesives, and insecticides (Glüge et al., 2020).

However, these unique characteristics also confer resistance to natural degradation of PFAS, impeding metabolism and excretion by organisms upon absorption and are known for its environmental persistence, biomagnification, and bioaccumulation (Death et al., 2021; Lu et al., 2023; Sharma et al., 2024). Humans are exposed to PFAS through ingestion of contaminated food, water, soil, and inhalation of contaminated air (Brunn et al., 2023; Cordner et al., 2019). Detectable PFAS level were found in most human populations, and in the United States, nearly all adults have some level of exposure. This high exposure persists despite efforts to reduce or eliminate production globally (ATSDR, 2023; Brunn et al., 2023; Gaines, 2023).

Once consumed, PFAS can induce a range of harmful outcomes, including carcinogenesis, genotoxicity, neurotoxicity, reproductive toxicity, developmental toxicity, and renal toxicity (Fenton et al., 2021; Hekster et al., 2003; Steindal and Grung, 2021). Concerns regarding PFAS exposure and its potential adverse effects on kidney health have escalated over time (Stanifer et al., 2018; Zhang et al., 2023). While the liver has traditionally been considered the primary target organ of PFAS toxicity, emerging evidence suggests that the kidney is also vulnerable to PFAS-induced damage (Liu et al., 2023; Rashid et al., 2020b; Solan et al., 2023; Stanifer et al., 2018; Zhang et al., 2023). In a 30-day exposure experiment in rats, the cumulative concentration of PFOA in the kidney was found to be the highest at 228 ± 37 μg/g, highlighting the organ’s vulnerability. (Cui et al., 2009). Our previous work has shown that PFOA and PFOS leads to altered gene expression in apoptosis and key metabolisms, altered DNA methylation patterns (Liu et al., 2023; Rashid et al., 2020b; Wen et al., 2022b). Given the array of new compounds in relation to its function, the relationship between PFAS exposure and kidney health is still under explored.

The emergence of energy-based PFAS such as lithium bis(trifluoromethanesulfonyl)imide (LiTFSI, known as HQ-115), an electrolyte salt used in lithium-ion batteries, further amplifies these concerns. Despite its detection in surface and wastewater near manufacturing sites, limited toxicity data is available for LiTFSI (EPA., 2023). Given the growing demand for rechargeable lithium batteries and the low battery recycle rate, the world faces the prospect of widespread release of PFAS such as LiTFSI and its counterparts (Guelfo et al., 2023) and hence the heightened concern.

Recent years have witnessed efforts to elucidate the interplay between environmental toxicants and altered epigenetic mechanisms, particularly DNA methylation (Bollati and Baccarelli, 2010; Ho et al., 2012). PFAS, classified as endocrine-disrupting chemicals (EDCs), have the capacity to mimic endogenous hormones, affecting gene expression leading to adverse health effects through altered epigenetic programming (Crews and McLachlan, 2006; Jacobs et al., 2017). In our study, we aim to elucidate the potential nephrotoxicity of LiTFSI, investigate epigenetic alterations using Reduced-representation bisulfite sequencing (RRBS), and explore the underlying mechanisms of LiTFSI-induced nephrotoxicity. Comparison with PFOA exposure using identical dosing and exposure durations is provided as relevant.

## 2. MATERIALS AND METHODS

### Animal studies

Mouse exposure experiments were conducted as previously described (Sands et al., 2024). Fifty male CD1 mice were randomly assigned to ten groups, with five mice per group. Dosing of LiTFSI and PFOA (utilized as a positive control because this is one of the two most studied PFAS) began via oral gavage once daily when mice reached the age of 6 weeks and acclimated to their environment. The study utilized doses of 1 and 5 mg/kg/day for a 30-day exposure period and 10 and 20 mg/kg/day for a 14-day exposure period, all administered in corn oil or corn oil alone (as a vehicle control). The dose selection of LiTFSI was guided by the derivation of a noncancer 30-day reference value for oral exposure by the United States Environmental Protection Agency (EPA., 2023). PFOA doses were based on human PFAS exposure ranges outlined by the Agency for Toxic Substances and Disease Registry (ATSDR, 2023), encompassing exposure scenarios in the general population, contaminated communities, and occupational settings. Mice had ad libitum access to Teklad Rodent Diet 8604 and purified water throughout the experiment. Following the exposure periods, mice were humanely euthanized via CO2 asphyxiation followed by cervical dislocation. Kidneys were promptly excised, weighed, and stored at -80°C. Relative kidney weight was calculated using the formula: Relative kidney weight = (Kidney weight / Body weight) * 100. All experiments were conducted in accordance with an approved protocol (Toxicology of Endocrine Disrupting Chemicals, Protocol # 22038) by the University of Illinois Urbana-Champaign Institutional Animal Care and Use Committee (IACUC) in compliance with National Institute of Health (NIH) guidelines.

### Biochemical measurements

Blood was promptly drawn from their hearts after CO2 asphyxiation and kept on ice until centrifugation at 4000 rpm for 15 min to isolate serum. The serum was then stored at -80°C prior to conducting biochemical measurements. Serum levels of various biochemical parameters, including Blood urea nitrogen (BUN), Creatinine, Bicarbonate, were analyzed using a Beckman Coulter DxC700 AU at the Veterinary Diagnostic Laboratory, University of Illinois Urbana-Champaign.

### LiTFSI & PFOA content in serum and kidney

LiTFSI (HQ-115) and PFOA were isolated from serum and kidney tissues using our previously established method (Wen et al., 2022a). The concentrations of LiTFSI and PFOA were determined using liquid chromatography-mass spectrometry (LC-MS), employing both internal and external standards, with calibration conducted across various concentration levels. LC-MS analyses were conducted by the Carver Metabolomics Core Facility at the Roy J. Carver Biotechnology Center, University of Illinois Urbana-Champaign, utilizing an Agilent 1260 Infinity II HPLC system (Agilent Technologies, Santa Clara, CA, USA). Metabolites were separated on a Phenomenex Kinetex PS C18 100A column (2.6 µm, 100 x 4.6 mm) (Phenomenex, Torrance, CA, USA) using a gradient method with mobile phase A (H2O + 20 mM ammonium acetate) and mobile phase B (methanol) at a flow rate of 0.35 mL/min. The gradient profile was as follows: 0-2 min = 90% A; 2-10 min = 0% A; 18.1-24 min = 90% A. Detection was performed using a Sciex 6500+ triple quadrupole MS (Sciex, Framingham, MA, USA) operating in the negative ionization mode.

### Histological staining

After the exposure period and euthanasia, fresh kidney samples from mice treated with LiTFSI and PFOA were promptly immersed in 10% neutral buffered formalin (Sigma-Aldrich; St. Louis, MO, USA) for 24 hours at room temperature. Subsequently, they were transferred to 70% ethanol solutions. Following dehydration, the kidneys were embedded in paraffin and sliced into sections 5-μm thick. Hematoxylin and eosin (H&E) staining (Sigma-Aldrich, St. Louis, MO, USA) was performed on the sectioned slides, and images were captured using a brightfield microscope.

### gDNA extraction

Kidney tissues were harvested from the following groups: mice exposed to 20 mg/kg/day of LiTFSI/PFOA for 14 days, mice exposed to 5 mg/kg/day of LiTFSI/PFOA 30-dayally, vehicle control groups for both 14-day and 30-day exposures. Genomic DNA extraction and purification were performed on these samples using the Purelink genomic DNA mini kit (Thermofisher, Waltham, MA, USA) following the manufacturer’s instructions. RNase A treatment was conducted to remove RNA, as recommended by the manufacturer and necessary for subsequent RRBS analysis. The concentration of extracted DNA was determined using a Qubit assay, and its quality was evaluated using the Fragment analyzer and DNA electrophoresis gel.

### Library construction and sequencing by Novaseq 6000

RRBS library construction and sequencing were performed at the Roy J. Carver Biotechnology Center at the University of Illinois at Urbana Champaign using Illumina Novaseq 6000. Libraries were constructed with the Ovation RRBS Methyl-Seq kit from Tecan, CA. Briefly, 100 ng of high molecular weight DNA was digested with MspI enzyme, ligated to sequencing adaptors, treated with bisulfite, and amplified by PCR. The final library concentrations were quantified with Qubit (ThermoFisher, MA) and the average size was determined with a Fragment Analyzer (Agilent, CA). The libraries were then diluted to 10 nM and further quantitated by qPCR on a CFX Connect Real-Time qPCR system (Biorad, Hercules, CA) for accurate pooling of barcoded libraries and maximization of the number of clusters in the flow cell.

The pooled barcoded shotgun libraries were then loaded on a NovaSeq 6000 SP lane for cluster formation and sequencing. They were sequenced for 100 nucleotides from one side of the DNA fragments. The typical output per lane in the NovaSeq is 400 million reads (SP flowcell) and we obtained approximately 30 million reads per sample. The FASTQ read files were generated and demultiplexed and adapters were trimmed with the bcl2fastq v2.20 Conversion Software (Illumina, San Diego, CA).

### RRBS Data analysis

Following RRBS analysis of the LiTFSI treatment group (Vehicle control: n = 3; 20mg/kg/day LiTFSI under 14-day exposure: n=3; 5mg/kg/day LiTFSI under 30-day exposure) and the PFOA treatment group (control: n = 3; 20mg/kg/day PFOA under 14-day exposure: n=3; 5mg/kg/day PFOA under 30-day exposure), we employed the modified Nf-core methylseq pipeline (Ewels et al., 2020) (10.5281/zenodo.2555454) for sequence alignment and methylation calls extraction (Nextflow version: 22.09.7.edge). Trimmed reads were aligned to the mouse reference genome GRCm39, excluding the first four base pairs due to adapter contamination. The deduplication step was omitted for this analysis. Alignment quality was assessed using FastQC, and methylation calling was conducted using Bismark version 0.22.4 (Krueger and Andrews, 2011). Known C-T SNPs, identified as a potential error source in RRBS data analysis, were determined using merged sorted BAM files generated after alignment through BS-SNPer (Gao et al., 2015) with default settings (10x coverage was used to align with the parameters in downstream analysis).

Extracted methylation calls served as inputs for methyKit (version 1.27.1) (Akalin et al., 2012) in R (version 4.3.1). Reads with a minimum 2x coverage were imported using methylKit::methRead. Samples were filtered by discarding coverage below ten reads and bases exceeding the 99.9th percentile. Sample data were normalized using the default "median" method. C-T SNPs were removed as a further filtering step. After filtering and normalization, CpGs covered in all libraries were retained for exploratory analysis and examination of the global methylation distribution.

Differential DNA methylation was calculated by comparing the proportion of methylated Cs in control samples versus samples with 14-day and 30-day exposure. The logistic regression test was employed for differential DNA methylation calculation. Resulting p-values were automatically generated by multiple testing using the Benjamini-Hochberg FDR method. Differentially methylated CpGs (DMCs) were defined as CpG sites not overlapping with C-T SNPs, exhibiting a percentage methylation difference >50 and a q value <0.01. The type was specified as "hyper" or "hypo" during DMC extraction. The locations of DMCs were summarized in pie charts. Differentially methylated regions (DMRs) were defined using a tiling window of 100bp resolution, including the above CpGs with at least a 25% difference in methylation and a q value <0.01.

### KEGG and GO enrichment

Gene ontology (GO) and Kyoto Encyclopedia of Genes and Genomes (KEGG) represent widely adopted methods for functional analysis. Over Representation Analysis was employed to discern whether specific biological processes were enriched in genes based on differentially methylated regions. To categorize genes with differentially methylated CpGs (DMC) into KEGG pathways or gene ontology for functional categorization, we utilized ClusterProfiler (Yu et al., 2012; Wu et al., 2021). Here, both the P value cut-off and q value cut-off were set to 0.1. Figures presented in the results showcase the top ten categories.

### qPCR

Total RNA was extracted utilizing the Quick-RNA Miniprep Kit (R1054, Zymo Research), which effectively removes genomic DNA. The quantity and quality of the isolated RNA were evaluated using NanoDrop. Subsequently, cDNA synthesis was conducted using the high-capacity cDNA Reverse Transcription Kit (Applied Biosystems REF 4368814). Analysis of mRNA was performed via quantitative reverse transcription polymerase chain reaction (qRT-PCR), utilizing SYBR Green PCR Master Mix reagents (Applied Biosystems REF 4367659) following the manufacturer’s instructions, and the Step One Plus Real-Time PCR System (Applied Biosystems). Relative quantification was achieved by normalizing to the expression levels of Beta-2-Microglobulin (*B2m*) and TATA-Box Binding Protein (*Tbp*), with reactions carried out in triplicate. The mRNA expression levels were determined using the 2-ΔΔCt method for relative quantification of gene expression (Taylor et al., 2019). All primer sequences are listed in **Table S6**.

### Statistical analysis

The data were analyzed using SPSS software, and graphical representations were generated using GraphPad Prism 9 Software. Students’ t-test was employed to assess the statistical significance of differences between two experimental groups. Analysis of variance (ANOVA) was utilized to evaluate potential statistically significant differences among the means of multiple treatment groups. A significance level of P < 0.05 was considered indicative of statistical significance.

## 3. RESULTS AND DISCUSSION

### LiTFSI exposure induced alterations in kidney-related biochemical indicators in male mice

Relative kidney weight and various biochemical parameters were assessed to evaluate renal function. Our findings revealed that LiTFSI exposure, at doses of 10 and 20 mg/kg/day under 14-day exposure and 1 and 5 mg/kg/day under 30-day exposure, did not significantly affect serum blood urea nitrogen (BUN) and serum creatinine levels (**Figure S1 e-f**). However, a significant decrease in serum bicarbonate levels was observed at doses of 10 and 20 mg/kg/day under 14-day exposure (**Figure 2C**). There were no significant alterations in body weights or relative kidney weights in LiTFSI-exposed male mice (**Figure S1 a-d**). Additionally, evaluation of PFOA-exposed mice revealed significant reduction in body weight (**Figure S1 a-b**), but no significant changes were observed in relative kidney weight (**Figure S1 c-d**). Moreover, a decrease in serum BUN levels was observed in the 20 mg/kg/day group under 14-day exposure, along with decreased serum bicarbonate levels under 14-day treatment (**Figure 2A** and **Figure 2C**). These results suggest that LiTFSI exposure altered kidney-related biochemical indicators without influencing relative kidney weight. Our findings are consistent with previous studies which show that exposure to PFAS can induce kidney injury, resulting in altered levels of kidney injury markers, such as creatinine, uric acid, and BUN (Hall et al., 2023; Jain and Ducatman, 2019a; Jain and Ducatman, 2019b). A significant decrease in serum bicarbonate levels may indicate potential kidney injuries, as evidenced by previous studies linking reduced bicarbonate levels with chronic kidney disease (CDK) in many studies (Menon et al., 2010; Raphael et al., 2014; Rashid et al., 2020b; Shah et al., 2009).

**Figure 1.**
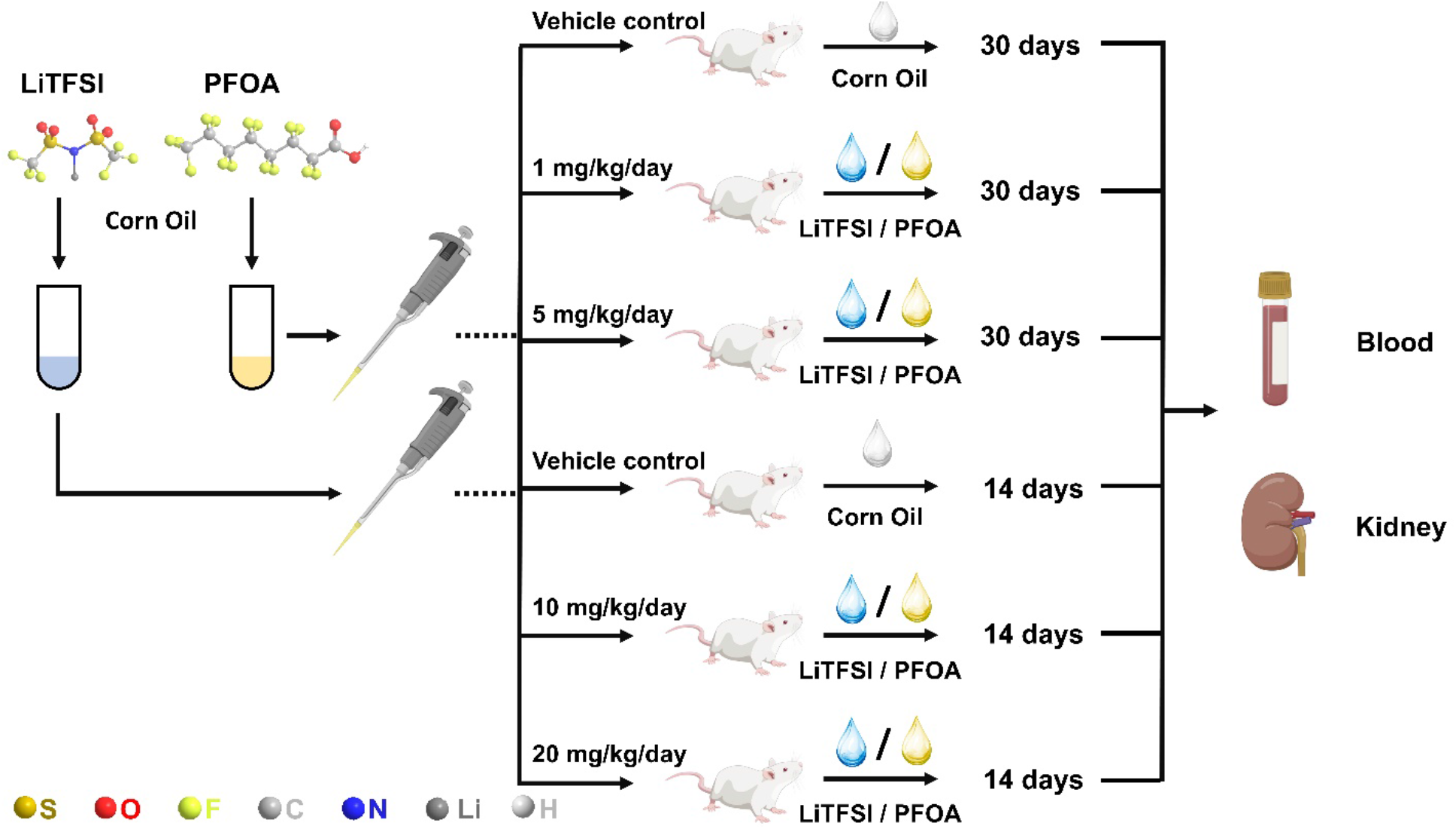
Overview of animal exposure design, featuring two dosage levels administered over 30 days (1 and 5 mg/kg/day) and two dosage levels administered over 14 days (10 and 20 mg/kg/day).

**Figure 2.**
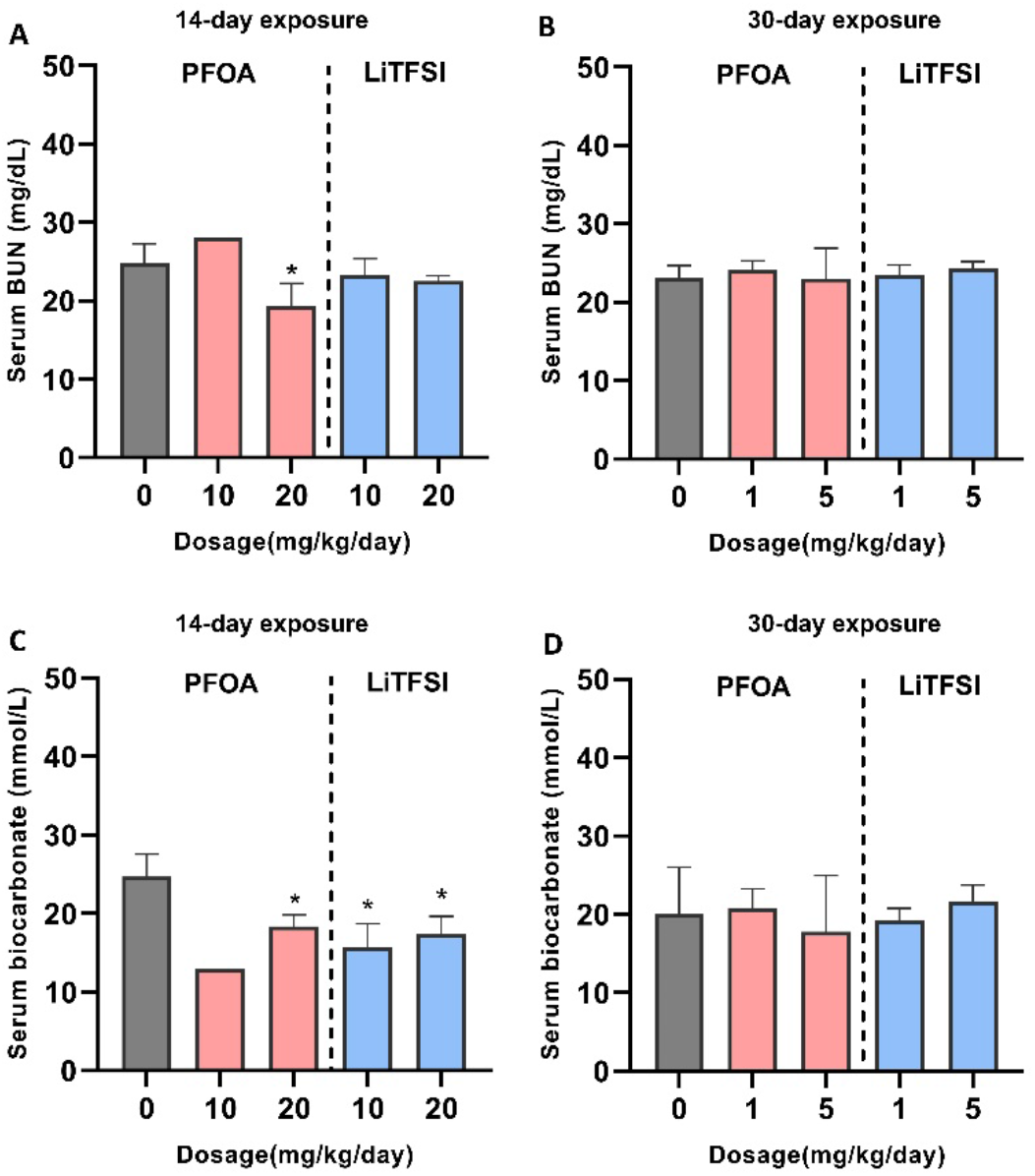
Effects of LiTFSI and PFOA on serum biochemical indicators in male mice. (A-B) Levels of Serum BUN. (C-D) Levels of Serum Bicarbonate. (n=4 per group). Data are presented as mean ± SD. *P< 0.05.

### LiTFSI exposure induced kidney injuries and inflammatory response in male mice

Histological changes were evaluated to elucidate the renal injuries induced due to exposure to LiTFSI. Kidney tissue sections stained with H&E revealed normal renal structures in the vehicle control groups (**Figure 3A** and **Figure 3D**), with no evident damage observed. In contrast, various degrees of histological changes were evident in both LiTFSI and PFOA-exposed groups. The LiTFSI 14-day exposure group (20 mg/kg/day) exhibited tubule dilation, irregular tubular shape, and proliferation in renal corpuscles (**Figure 3B**). More severe kidney damage was observed in the LiTFSI 30-day exposure group (5 mg/kg/day), characterized by inflammation, congestion, tubule dilation, loss of kidney structure, and proliferation of renal corpuscles (**Figure 3C**). In the PFOA-treated groups, inflammation, congestion, and proliferation in renal corpuscles were detected, with severe damage observed in the groups with longer exposure periods (**Figure 3C** and **Figure 3F**).

**Figure 3.**
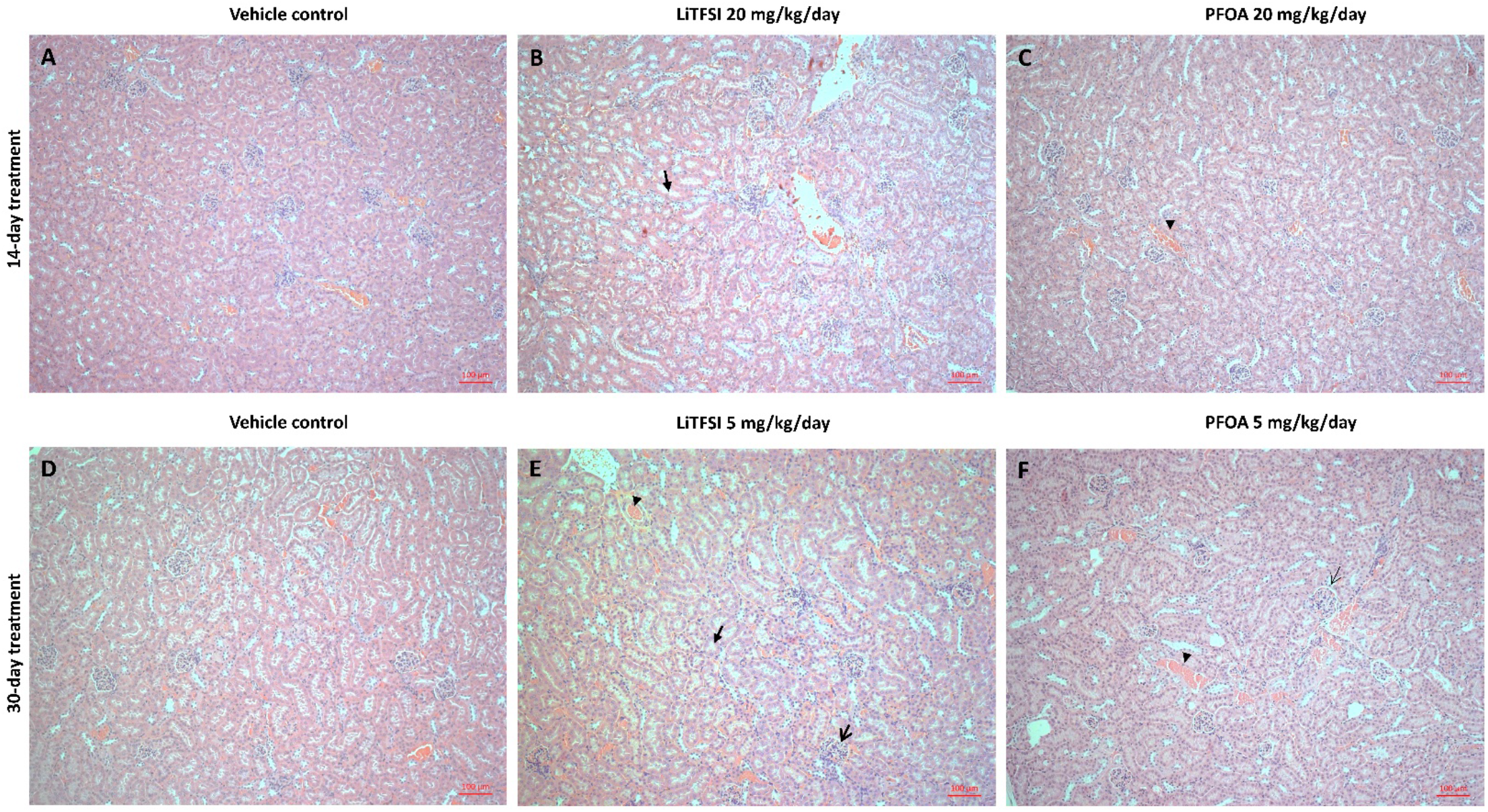
Effects of LiTFSI and PFOA on kidney histology in male mice. (n=3 per group). Renal tubule dilation (filled arrow) was observed in LiTFSI treatments. Proliferation in renal corpuscles (open arrow) were observed in both LiTFSI and PFOA treatments. Inflammation and congestion (Open arrow) were observed in PFOA treated and 30-day LiTFSI treated groups.

Major kidney injury markers IL-6, TNF-α, TGF-β, PPARα, and IFN-γ (Anders et al., 2010; DONNAHOO et al., 2000; Iwaki et al., 2019a; Lan and Chung, 2012; Nechemia-Arbely et al., 2008) were evaluated. A significant increase in TNF-α gene expression was observed in the 14-day LiTFSI treated groups, indicating an inflammatory response. Conversely, significant decreases were observed in TGF-β and PPARα genes under 30-day LiTFSI treatment (**Figure 4**). These findings suggest that subchronic and subacute exposures may lead to different regulatory responses. In the PFOA-exposed groups, we noted a consistent elevation in all five kidney injury markers across both 14-day and 30-day exposure periods. This observation is aligned with previous studies on the deleterious effect of PFOA and its ability to induce kidney injuries (Cui et al., 2009; Gong et al., 2019; Li et al., 2017; Rashid et al., 2020b).

**Figure 4.**
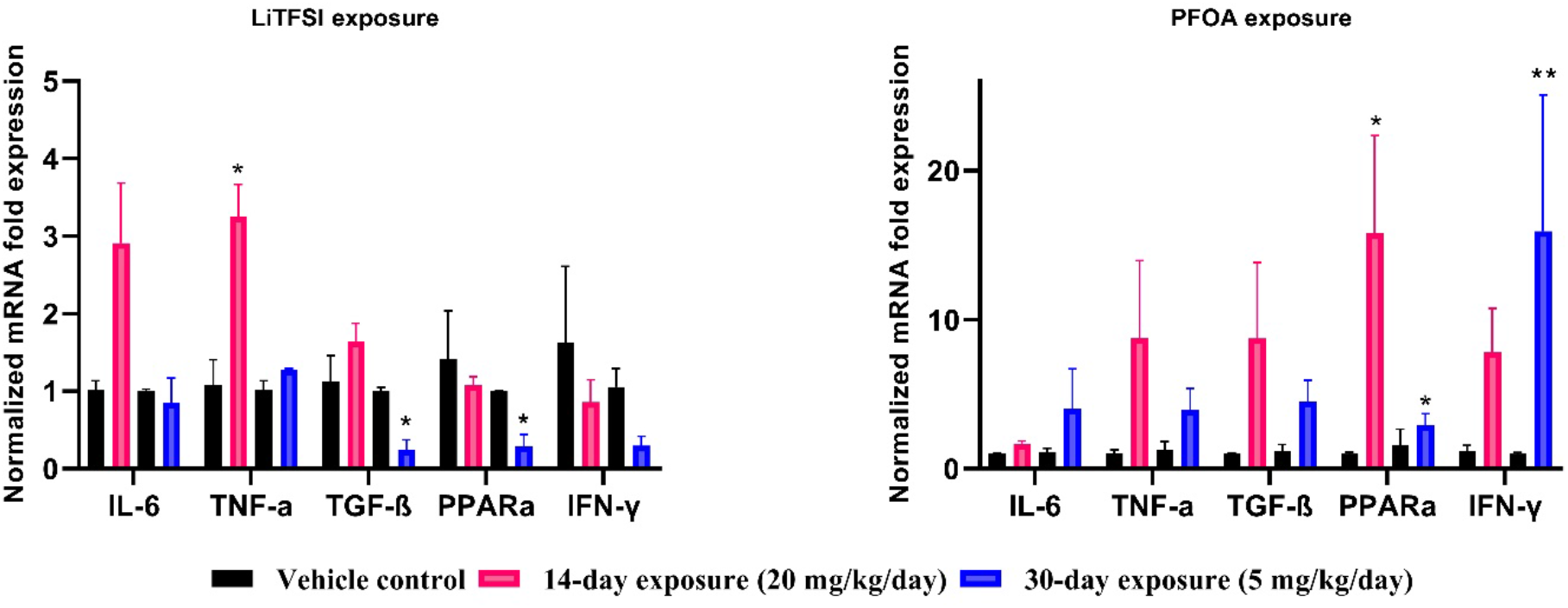
mRNA expression of five kidney injury markers was evaluated in LiTFSI/PFOA-exposed group (n=3 per group). Data are presented as mean ± SEM. ** P<0.01; *P < 0.05.

TNF-α, a proinflammatory cytokine, is implicated in renal injury by modulating hemodynamic and excretory functions in the kidney (Mehaffey and Majid, 2017; Ramseyer and Garvin, 2013). The elevated TNF-α expression observed under 14-day LiTFSI exposure indicates the presence of an inflammatory response. Transforming growth factor-beta (TGF-β) plays a dual role in immune response regulation. While known for its pro-fibrotic and pro-inflammatory properties, TGF-β also exhibits anti-inflammatory effects. By downregulating inflammatory processes, particularly in renal inflammation, TGF-β aids in reducing inflammation and promoting tissue healing and repair in the kidneys (Gu et al., 2020). The decrease in TGF-β expression in our study suggests a heightened inflammatory response. PPARα, a member of the nuclear hormone receptor superfamily, functions as a ligand-activated transcription factor. A reduction in PPARα gene expression may signify compromised kidney function and increased inflammation (Fruchart et al., 2020; Gao and Gu, 2022; Iwaki et al., 2019b). Overall, our histological and gene expression analyses reveal renal injuries and inflammatory response under LiTFSI exposure.

### Quality Control and Consistent Global DNA Methylation Patterns with DMR Methylation Profiles

To assess the impact of 14-day and 30-day exposure of LiTFSI on DNA methylation patterns in the kidney, we generated a total of 12 methylomes from male CD1 mice exposed to the contaminant (**Table S1**). For comparative analysis, PFOA-exposed groups were utilized as a relative control, since this is one of the two most studied PFAS, allowing us to discern the altered methylation profiles of a typical PFAS compound (PFOA) and the new LiTFSI. Employing RRBS, we obtained an average of 2.53 x 10^8^ paired end reads across eighteen libraries, post-adapter trimming, and quality control. A noteworthy 68.8% of the reads were uniquely aligned to the mouse reference genome GRCm39 (**Table S2**). To ensure the accuracy of global methylation patterns and downstream analysis, C-T single nucleotide polymorphisms (SNPs) were excluded. Each sample revealed approximately 1.41 x 10^6^ CpG sites (**Table S2**). We further explored the percentage of DMCs per chromosome, observing DMLs across all major chromosomes in both treatment groups (**Figure S2**).

Differential methylation at each base was computed using a logistic regression-based method, filtered by a q-value significance level of 0.01 and a percentage methylation difference exceeding 50%. In the LiTFSI treatment group, the 14-day exposure revealed that most CpGs (1,110,391) were either lowly or mildly methylated, with only 0.008% (88) displaying differential methylation. In the 30-day exposure group, 0.012% of CpGs (135) exhibited high methylation. Similarly, in the PFOA treatment group during 14-day exposure, the majority of CpGs (1,114,062) displayed low to mild methylation, with only 0.007% (82) showing high methylation. In the 30-day exposure group, 0.013% of CpGs (149) were highly methylated (**Figure 5A-D**; **Figure S2**). Subsequently, we identified DMRs by utilizing these CpGs and defining a 100bp sliding window with methylation levels exceeding 25% and a q-value below 0.01. In LiTFSI-exposed groups, we identified 128 DMRs in the 14-day exposure and 164 DMRs in the 30-day exposure groups. For PFOA, we found 157 and 181 DMRs in the 14-day and 30-day exposure groups respectively (**Figure 5E-H**). In summary, both LiTFSI and PFOA treatment groups displayed altered global and regional methylation patterns.

**Figure 5.**
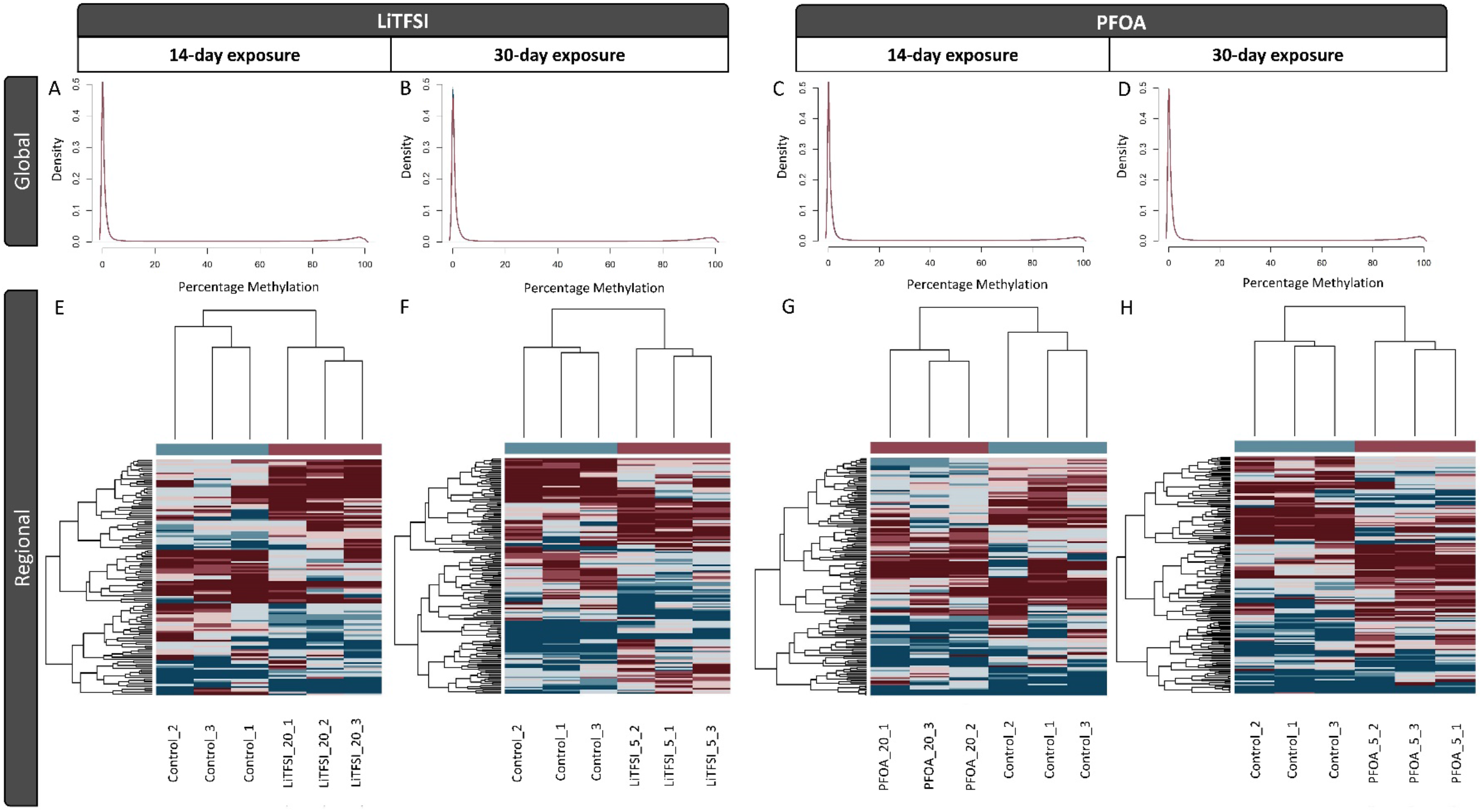
Global and regional DNA methylation profiles of kidney under LiTFSI. (A-D) Density plots of individual CpG methylation values. (E-H) Heatmaps of methylation changes for differentially methylated regions.

### Functional enrichment analysis implicate the involvement of different processes in the 14-day and 30-day exposure groups

Upon examination of the functional implications of LiTFSI exposure on developmental related genes, we conducted a comprehensive enrichment test analysis. This involved assessment of the biological relationships of genes mapped to DMRs in comparison with genes in their background regions. Notably, the top significant Gene Ontology enrichments in the 14-day exposure of LiTFSI were associated with biological processes such as DNA replication, mitochondrial transport, and smoothened signaling pathways, while molecular function terms included ATP-dependent activity and protein binding, among others. Conversely, in the 30-day exposure groups, biological processes were linked to cellular responses and transport, with molecular function terms encompassing hyaluronic acid binding, S-palmitoyl transferase activities, and S-acyltransferase activities (**Figure 6A**). Limited overlap was observed between the two exposure scenarios. Subsequently, our focus shifted to testing KEGG enrichments in both 14-day and 30-day LiTFSI exposures. The 14-day exposure showed top KEGG enrichments related to cellular senescence, PI3K-Akt Pathway, and p53 signaling pathways, while 30-day exposure featured enrichments related to Taste transduction, phospholipase D signaling pathway, non-homologous end joining pathways, and glucose metabolism (**Figure 6B**). We noted that there was no significant overlap between the pathways identified in the 14-day and 30-day exposure groups.

**Figure 6.**
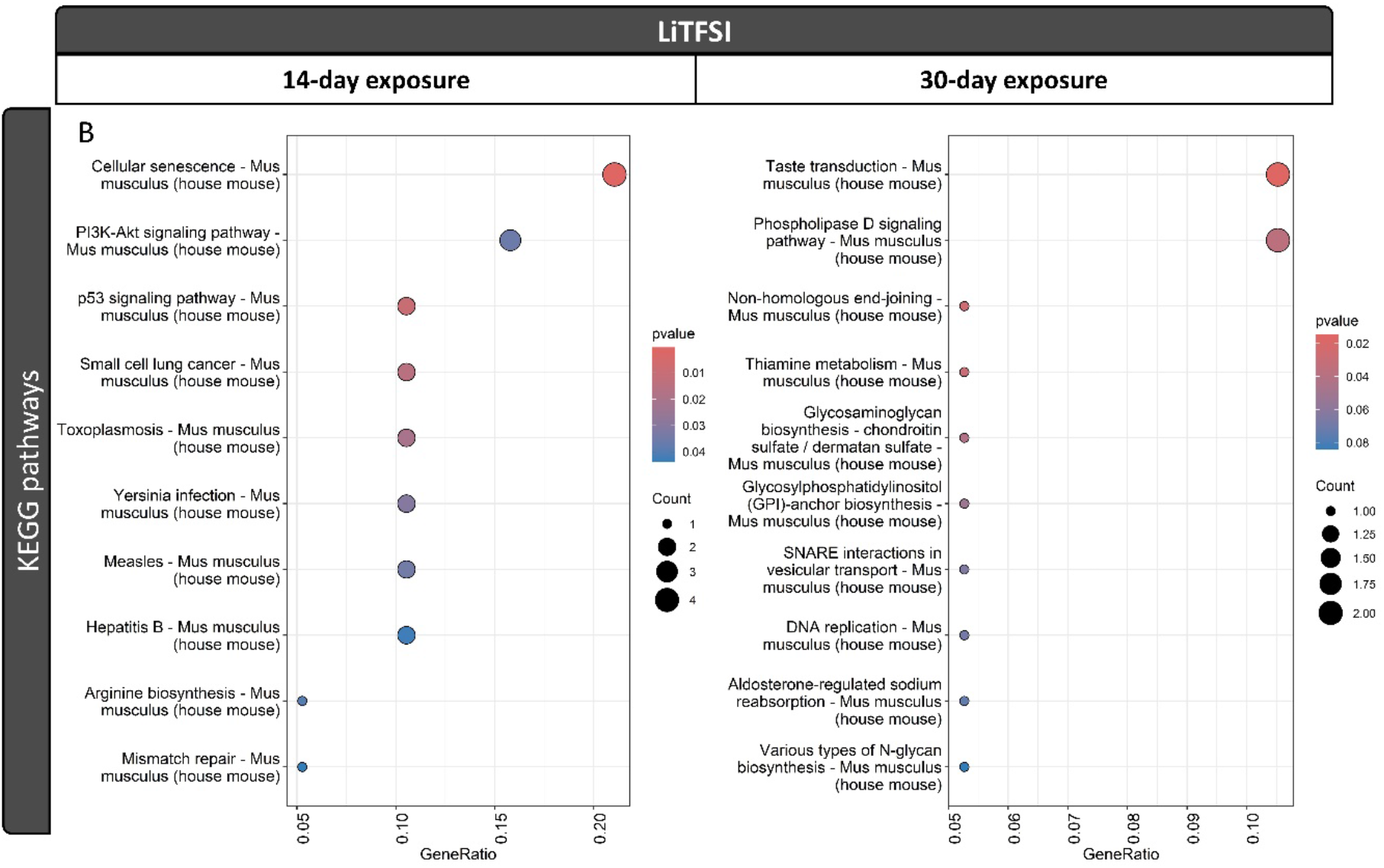
GO and KEGG enrichment results for kidney DMRs under LiTFSI treatments. (A) GO enrichment results (B) KEGG enrichment results.

In the 14-day LiTFSI exposed group, the top three KEGG pathways identified were cellular senescence, the PI3K-Akt Pathway, and the p53 signaling pathway. The PI3K-Akt Pathway is known to be frequently activated in various tumor types and is recognized as a cancer driver. Previous studies have demonstrated a high activation of the PI3K-Akt Pathway in renal cell carcinoma (RCC) (Guo et al., 2015; Park et al., 2007; Porta and Figlin, 2009). Furthermore, alterations in the cellular senescence-related p53 signaling pathway are also indicative of RCC (Amendolare et al., 2022; Fu et al., 2022; Higgins et al., 2018; Kurbegovic and Trudel, 2020). Our results suggest that methylation patterns in key genes within these pathways may contribute to kidney injury by promoting inflammation, abnormal cell proliferation, and apoptosis, warranting further focused studies.

Additionally, we explored gene enrichment in the PFOA-treatment group. In the 14-day exposure groups, significant GO enrichments were observed in molecular function terms such as ATP-dependent activity, protein binding, DNA helicase activity, and catalytic activity. Biological processes included endosomal toll-like receptor signaling pathways and cell surface toll-like signaling pathways. Conversely, in the 30-day exposure group, molecular function terms related to hyaluronic acid binding, and biological processes linked to glycosylation activities and glycolipid biosynthetic processes were prominent (**Figure S4A**). As observed in LiTFSI exposure groups, limited overlap was observed between the two PFOA exposure scenarios. KEGG pathway enrichment revealed distinct terms, with 14-day exposure featuring coronavirus disease and mismatch pair, while the 30-day exposure showed thiamine metabolism as the top significant term (**Figure S4B**). Again, minimal sharing of pathways was evident between the 14-day and 30-day exposure groups in PFOA-treated mice.

### Enrichment of LiTFSI Exposure DMRs in Intergenic Regions and development-related Motifs

To assess the hypothesis that LiTFSI exposure induces methylation variations linked to gene regulation, we conducted an enrichment analysis of LiTFSI and PFOA DMRs, focusing on annotated genomic regions compared to background regions. Initial scrutiny of CpG islands revealed that a majority of DMRs were enriched outside CpG islands and shores in both 30-day and 14-day exposure groups of LiTFSI and PFOA (**Figure 7**; **Table S3**). Specifically, LiTFSI DMRs and PFOA DMRs demonstrated significant enrichment within intergenic or intron regions (**Figure 7A & C**). Further investigation of LiTFSI DMRs aimed to ascertain their functional relevance in gene regulation relative to background regions. Specifically, we explored enrichment for known transcription factor binding motifs (**Figure 7B & D**). In LiTFSI 14-day-exposed DMRs, the most significantly enriched Homer *de novo* motif was Bcl11a, a development-related transcription factor widely expressed in hematopoietic systems. Additional enriched motifs included CTCFL, implicated in the regulation of the PI3K-Akt pathway; Hand1::Tcf3, part of the Tal-related::E2A family; NF1-halfsite; and Homer known motifs, circadian rhythm motifs (BMAL1 and CLOCK). In LiTFSI 30-day exposure, Homer *de novo* motifs such as ONECUT1, regulating liver-expressed genes; BHLH family motifs (Arnt:Ahr); TALE family motif (PKNOX2); and Homer known motif RUNX2, associated with bone development, were enriched in our DMRs.

**Figure 7.**
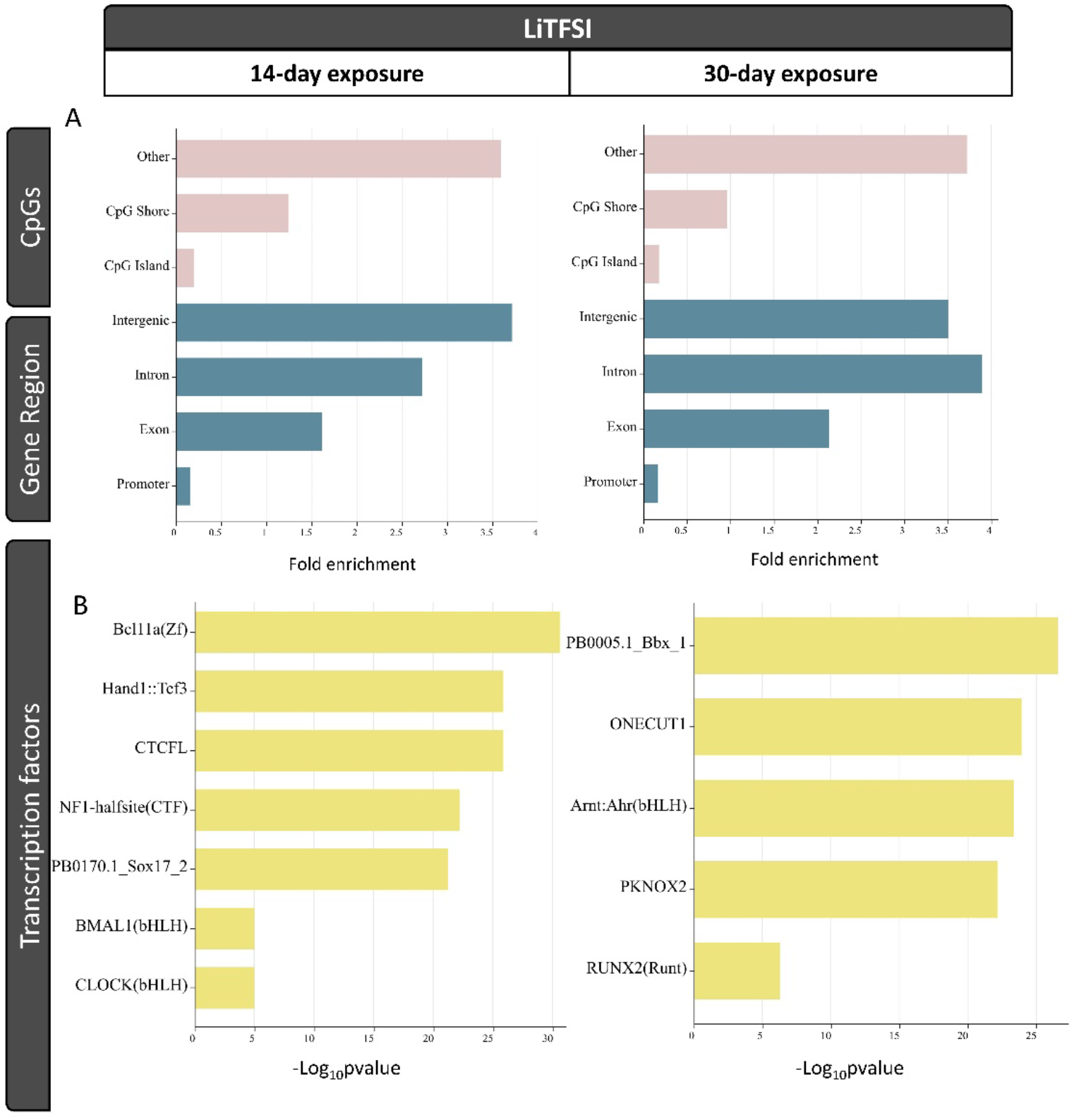
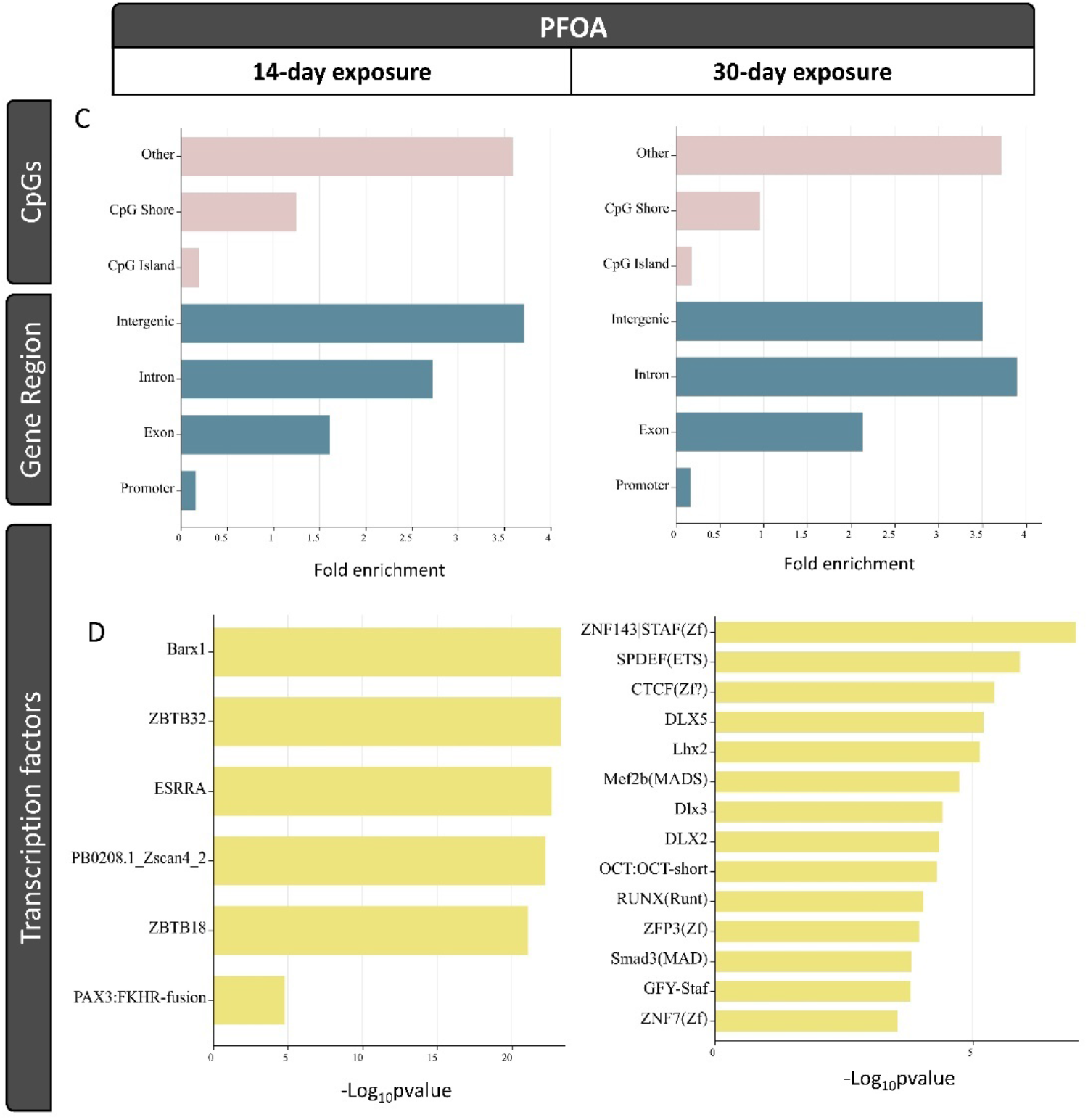
Annotation enrichment analysis for DMRs induced by LiTFSI and PFOA treatments in the kidney. (A & C) display CpG and gene region annotation enrichments. (B & D) present transcription factor enrichments. The significance threshold for p-values is set at e^-9^.

Upon examining the PFOA-14-day exposed group, enriched motifs included C2H2 zinc finger factors (ZBTB32, ZBTB18); BARX1, highly expressed in the stomach mesenchyme during gut organogenesis; and ESRRA, a key regulator of intestinal homeostasis through the activation of autophagic flux via gut microbiota. In 30-day PFOA-exposed DMRs, enriched motifs comprised ZNF family motifs (ZNF143, CTCF, ZNF3, ZNF7); ETS family motif (SPDEF), associated with tumor metastasis suppression; DLX motifs (DLX5, DLX2, DLX3), linked to oncogenesis; MADS-box motifs (Mef2b, Smad3), associated with various developmental processes; and LHX2, a regulator of neural systems.

Numerous studies have emphasized the pivotal role of CpG islands (CGIs) clustered in the promoter regions governing gene expression (Blackledge and Klose, 2011; Dhar et al., 2021; Hughes et al., 2020). However, our enrichment analysis revealed that genes associated with LiTFSI and PFOA DMRs were not significantly enriched in either promoter regions or CGIs, suggesting that DNA methylation might not be the sole contributor to kidney damage induced by LiTFSI or PFOA. Interestingly, we observed enrichment of development-related motifs in genes mapped to LiTFSI DMRs following both 14-day and 30-day exposures. This suggests that aberrant DNA methylation patterns may specifically occur in genes enriched with development-related motifs, potentially contributing to LiTFSI-induced kidney injury. For PFOA treatment groups, genes mapped to PFOA DMRs exhibited enrichment of motifs associated not only with development but also with carcinogenesis. This finding suggests that aberrant DNA methylation patterns may be linked to genes enriched with development-related motifs, contributing to PFOA-induced kidney injury.

### Genomic Coordination Overlaps Reveal Different Methylation Patterns in 14-day and 30-day Exposures but Consistent Trends Across Genders

Within the LiTFSI treatment group, approximately 9% of genomic coordinates were shared between 14-day and 30-day exposures, yet the majority exhibited differing methylation trends (**Figure 8A**). For instance, in Chr13:45023101-45023200, the mapped gene Jarid2 demonstrated hypermethylation under 14-day exposure but hypomethylation under 30-day exposure (**Table S4**). A similar phenomenon was observed in the PFOA treatment group (**Figure 8B**; **Table S4**). These findings suggest differential impacts of short-term and long-term exposures to LiTFSI and PFOA on DNA methylation patterns. Further exploration of the relationship between DNA methylation patterns and gene expression profiles (**Figure S5**) revealed no direct correlation between changes in methylation levels and gene expression. This indicates that alterations in DNA methylation may not solely dictate gene expression, and other epigenetic indicators such as the histone modifications may also play a role (Dong and Weng, 2013; Karlić et al., 2010). Additionally, factors such as the location of methylation and its interaction with transcription factors could influence gene regulation (Marchal and Miotto, 2015; Zhu et al., 2016).

**Figure 8.**
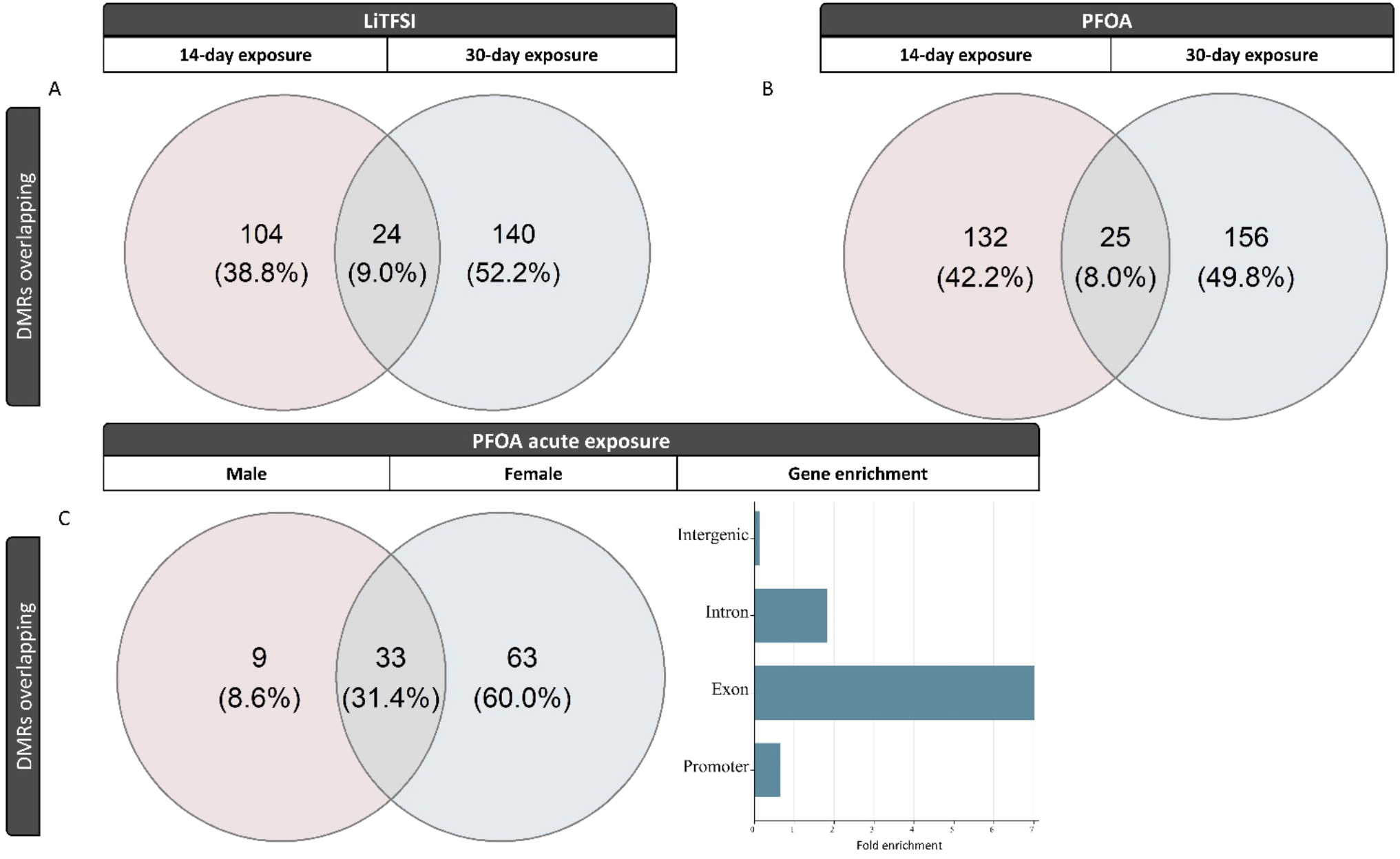
Genomic coordinates overlap in LiTFSI, PFOA-treated groups and in POFA 14-day treated male and 10-day treated female mice.

Furthermore, when comparing male mice exposed to PFOA for 14 days with our previously published data from female mice exposed to PFOA for 10 days (Rashid et al., 2020a), approximately 31.4% of genomic coordinates overlapped in both sexes (**Figure 8C**). Notably, despite the 14-day PFOA-treated male mice and 10-day PFOA-treated female mice leading to an enrichment of genomic coordinates in intergenic regions, the majority of overlapped genomic coordinates between the two sexes were primarily located within gene bodies and enriched in promoter regions (**Figure 8C**; **Table S5**). This underscores the influence of sex differences on PFAS-induced toxicity (Fenton et al., 2021).

### Effects of LiTFSI exposure on uric acid metabolism in male mice kidneys

The kidney plays a crucial role in regulating uric acid levels in the body by excreting excess uric acid. We further investigated whether exposure to LiTFSI affects uric acid metabolism in the kidneys. To understand the potential mechanisms involved, we examined the expression of several key genes involved in uric acid transport and regulation. Among the genes studied, ABCG2 is known as the urate exporter and expressed in the apical membrane of kidney proximal tubules, responsible for most of the extra-renal urate excretion (Huls et al., 2008; Ohashi et al., 2023). Additionally, we investigated organic anion transporter 1 (OAT1), which plays a role in urate secretion from the proximal tubules, facilitating the transport of urate from the cell to the interstitium (Nigam et al., 2007; Rouhani et al., 2018). GLUT9, also known as Glucose Transporter 9, is essential for regulating the reabsorption of uric acid from the urine back into the bloodstream in the kidney. (Auberson et al., 2018; Keembiyehetty et al., 2006). Another gene of interest was URAT1 (urate anion exchanger 1), responsible for urate reabsorption in the proximal tubules. (Shin et al., 2011). As shown in **Figure 9**, a significant decrease in GLUT9 and URAT1 expression was noted following 30-day exposure to LiTFSI. These transporters are primarily responsible for reabsorbing uric acid from the urine back into the bloodstream. The observed a decrease in their expression suggests a disruption in uric acid reabsorption, potentially leading to elevated uric acid levels in the urine and decreased levels in the bloodstream. Such alterations in uric acid metabolism may contribute to conditions such as hyperuricemia, a risk factor for various health issues. (Chung and Kim, 2021). Additionally, we observed changes in GLUT9 and URAT1 expression following 14-day exposure to PFOA, indicating PFOA also exerted effects on uric acid metabolism (**Figure 9**). These findings highlight the importance of understanding the impact of environmental exposures, such as LiTFSI and PFOA, on kidney function and uric acid metabolism, which could have implications on human health and disease risk. Further research should focus on elucidating the underlying mechanisms and potential health outcomes associated with altered uric acid metabolism due to environmental exposures.

**Figure 9.**
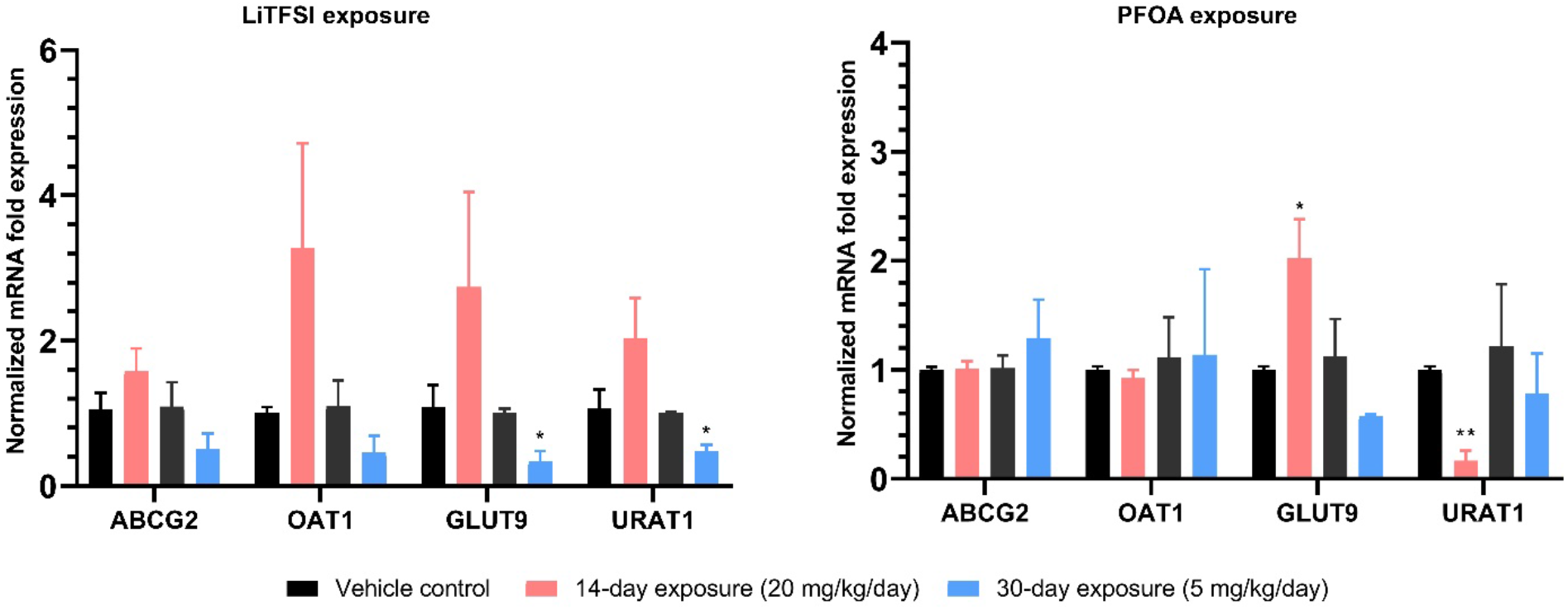
Effects of LiTFSI and PFOA on uric acid metabolism were evaluated (n=3 per group). Data are presented as mean ± SEM. ** P<0.01; *P < 0.05, as determined by non-parametric t-tests.

In summary, our investigation on the effects of LiTFSI exposure on kidney function, inflammatory responses, and epigenetic alterations in male mice sheds light on the potential health implications of this emerging energy-based environmental contaminant. We observed significant alterations in kidney-related biochemical parameters, histological changes indicative of kidney injuries, and dysregulated expression of genes associated with inflammation, renal function, and uric acid. Moreover, DNA methylation profiles revealed altered patterns in response to LiTFSI and PFOA exposure, with distinct functional enrichments in the 14-day and 30-day exposure groups. Enrichment of development-related motifs in LiTFSI DMRs and carcinogenesis-related motifs in PFOA DMRs suggests their involvement in kidney injury. Genomic coordination overlaps with the indicated differential methylation patterns between short-term and long-term exposure to LiTFSI and PFOA, with consistent trends across genders, highlighting the influence of sex differences on PFAS-induced toxicity. Additionally, LiTFSI exposure disrupted uric acid metabolism, evidenced by the decreased expression of key transporters involved in urate reabsorption. These findings underscore the nephrotoxic effects of LiTFSI and the importance of understanding the mechanisms underlying environmental contaminant-induced kidney dysfunction in the development of targeted interventions to mitigate adverse health effects.

## AUTHOR INFORMATION

### Author Contributions

M.S and X.Z contributed equally.

### Notes

Any additional relevant notes should be placed here.

## ACKNOWLEDGMENT

The authors would like to acknowledge Dr. Arnon gal for the assistance with histologic examination, the Roy J. Carver Biotechnology Center at the University of Illinois at Urbana-Champaign for assistance for sequencing support, the Carver Metabolomics core for assistance with LC-MS, and the Veterinary Diagnostic Laboratory for aiding in biochemical profiles. Biological assessments were performed at the Tumor Engineering and Phenotyping Facility in the Cancer Center at Illinois, while all animal-related procedures were carried out at the Beckman Institute.

