## Supplemental Information for "Kidney toxicology of a novel compound Lithium Bis(trifluoromethanesulfonyl)imide (LiTFSI, ie. HQ-115) used in energy applications: an Epigenetic evaluation"

### Contents list

Figure S1-S5 and Table S1-S6

Figure S1. Change in body weight, relative kidney weight, and serum creatinine levels following 14-day and 30-day exposure to PFOA and LiTFSI. (a-b) Body weight. (c-d) Relative kidney weight. (e-f) Serum creatinine levels. can do this when it comes for revision]

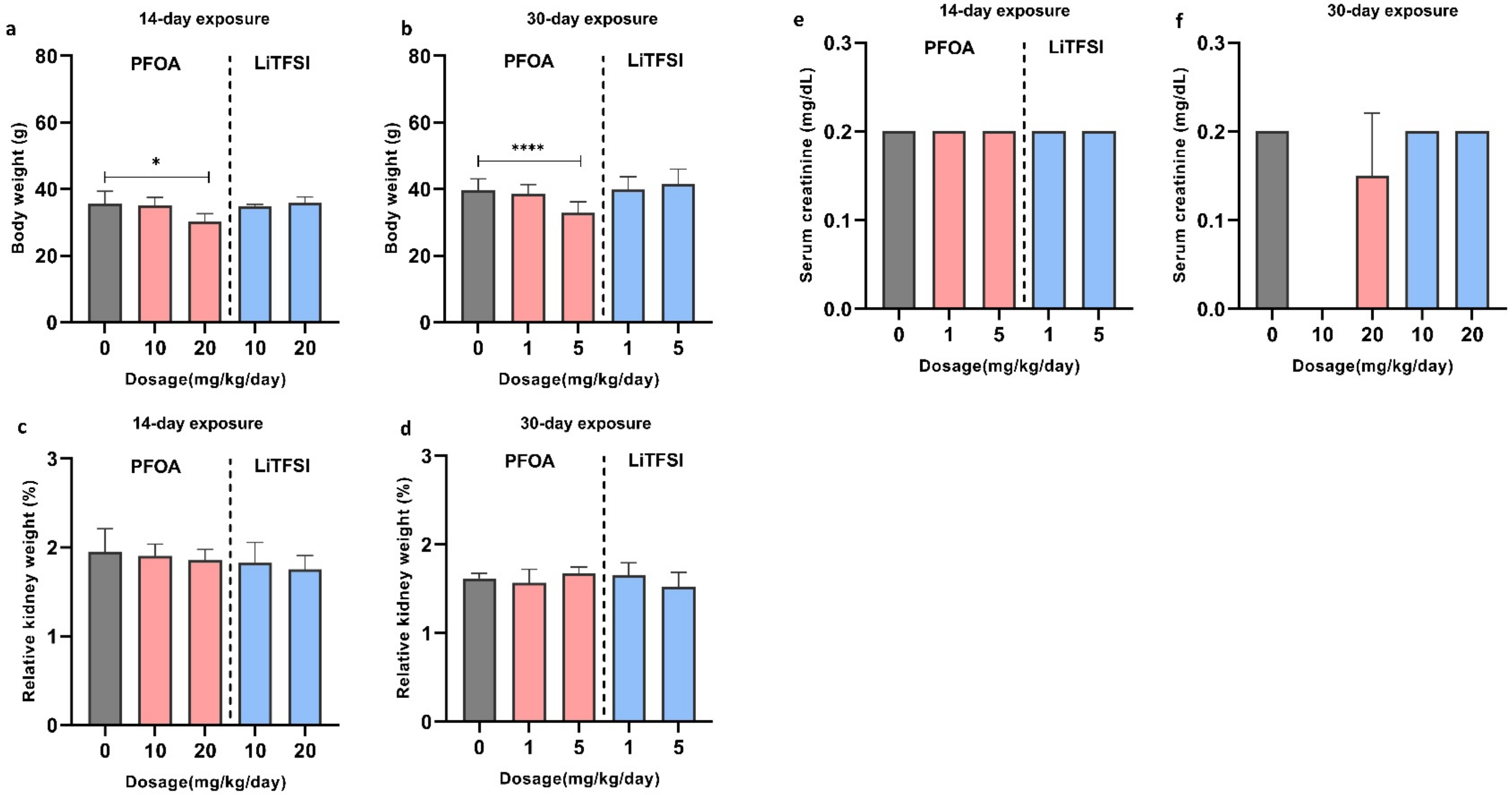

Figure S2. Global pattern of DMCs across all chromosomes.

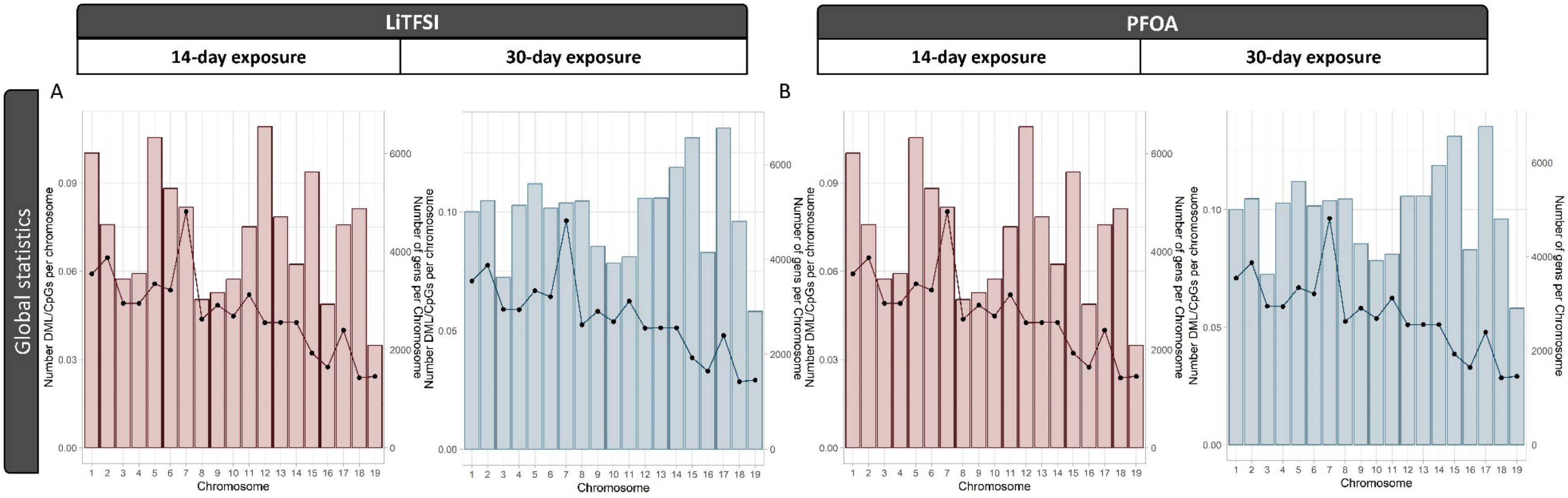

Figure S3. Global pattern of DMCs across all chromosomes.

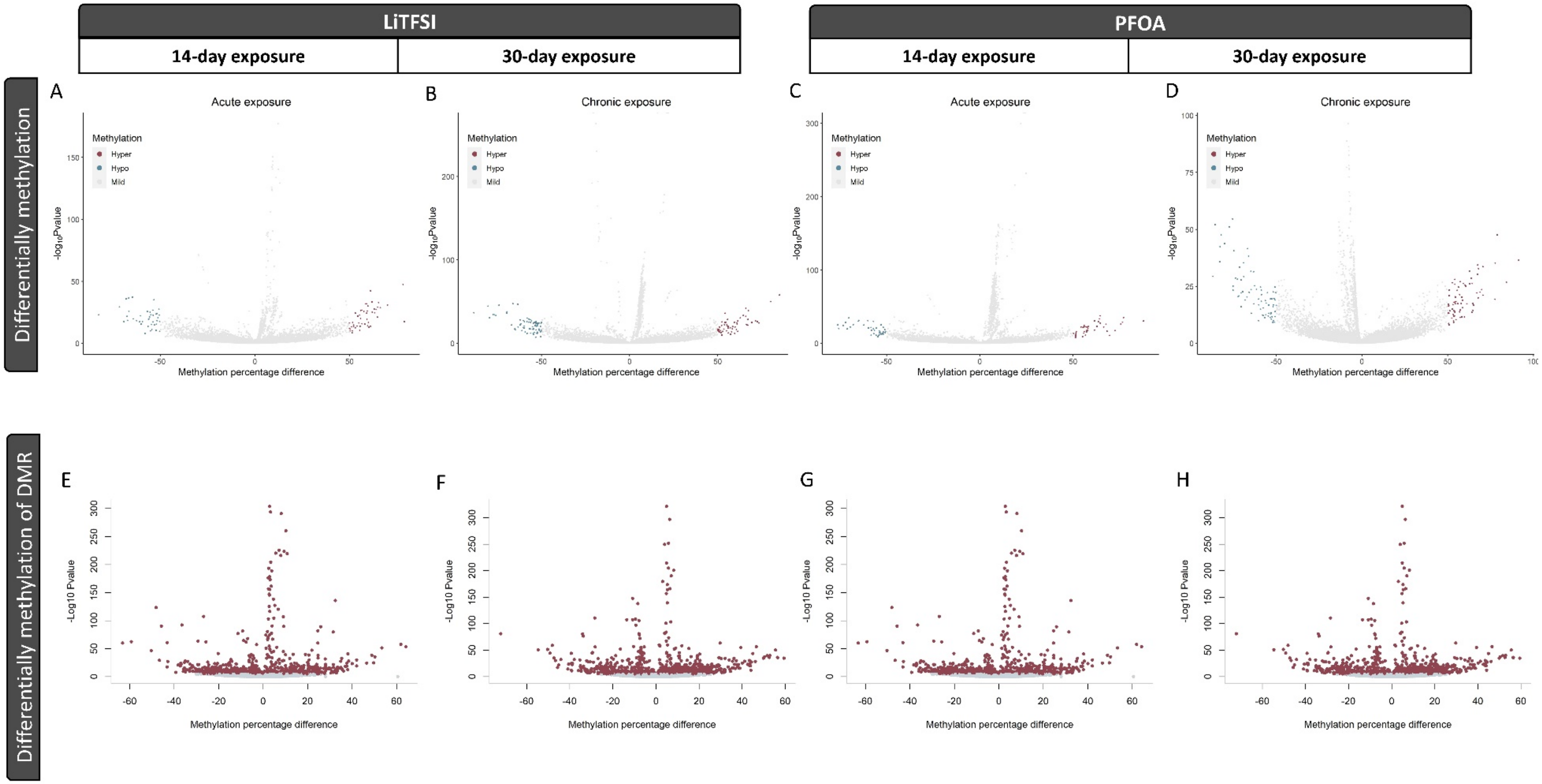

Figure S4. GO and KEGG enrichment results for kidney DMRs under PFOA treatment. (A) GO enrichment results (B) KEGG enrichment results.

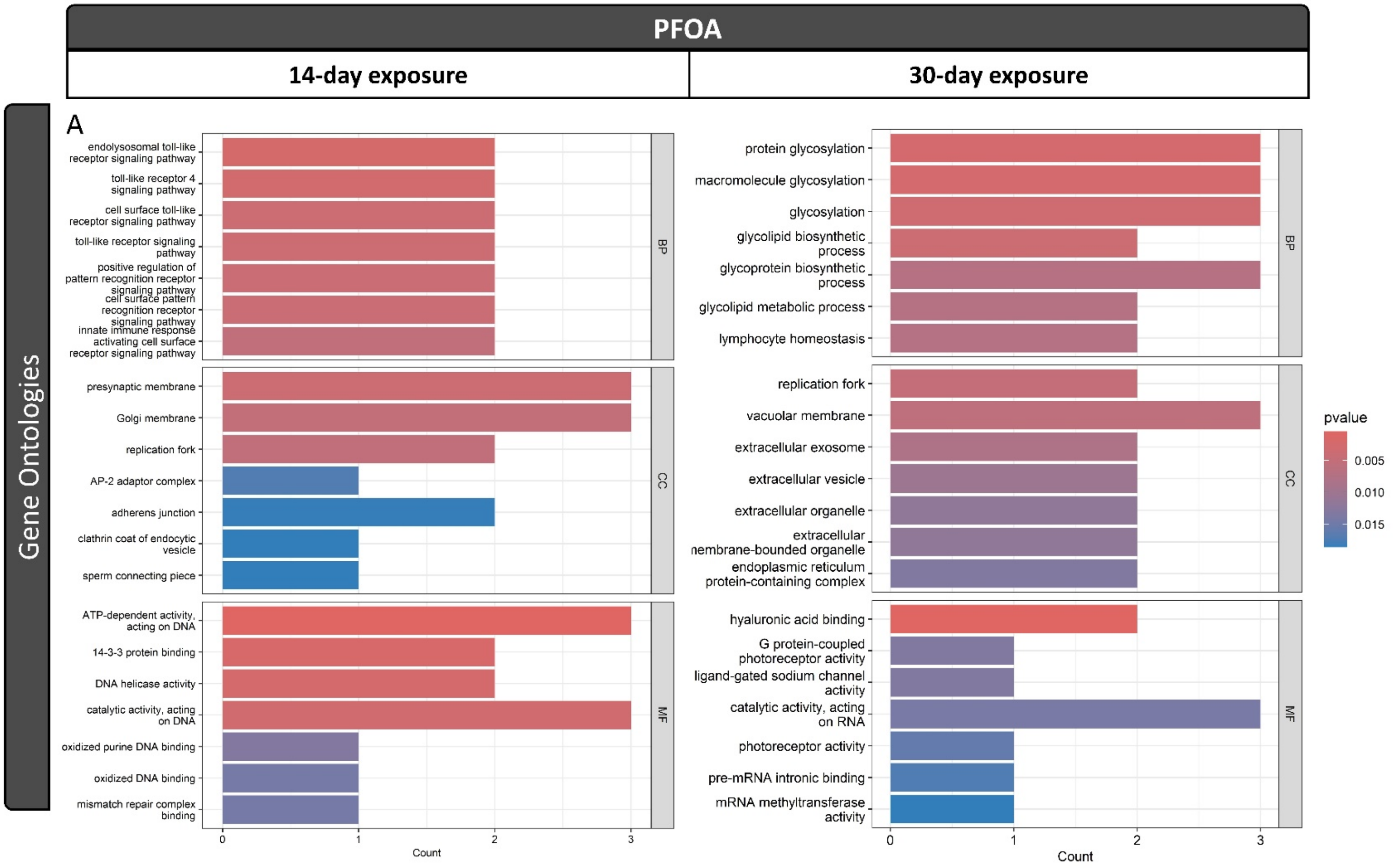

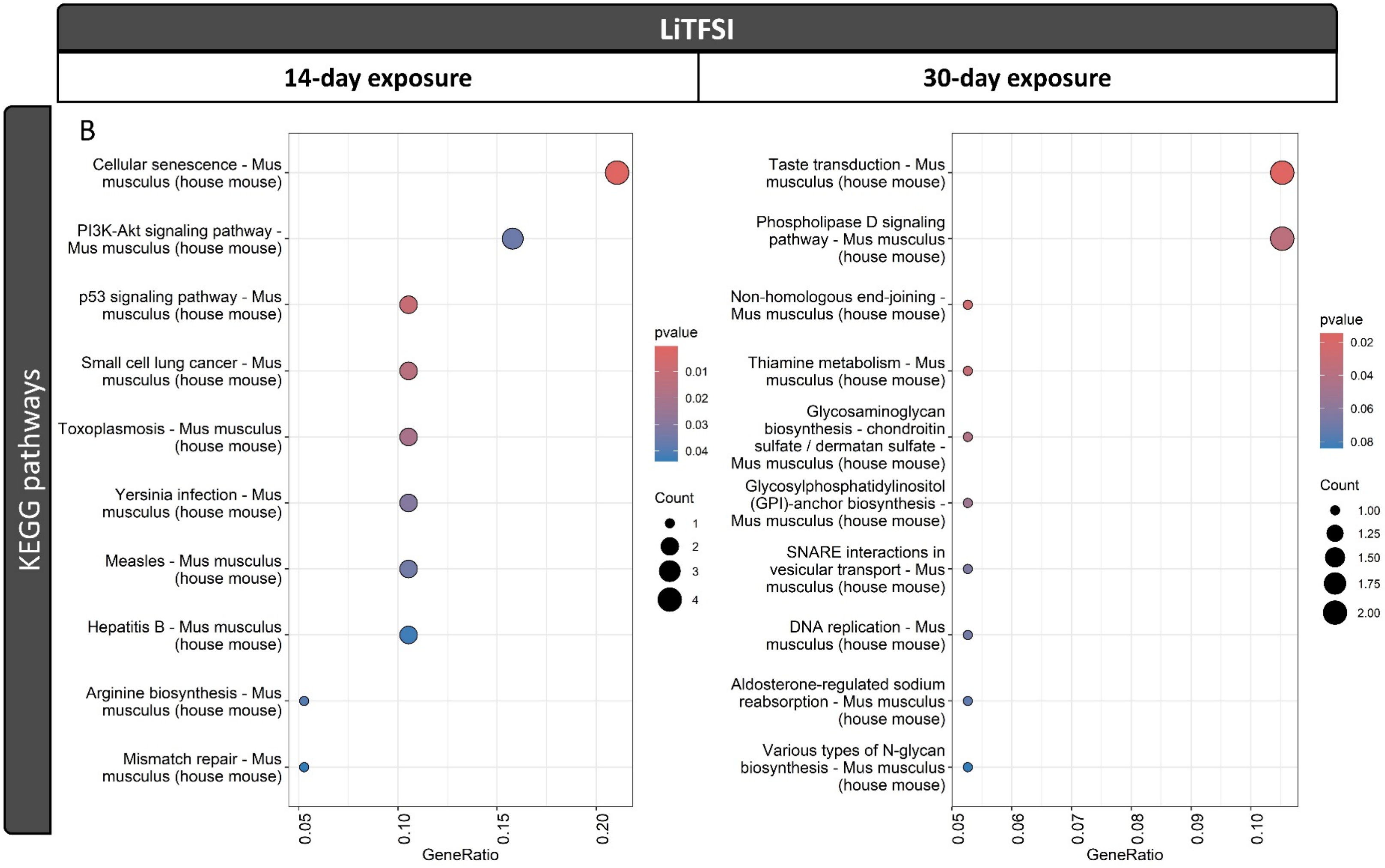

LiTFSI
14-day exposure
30-day exposure
KEGG pathways
B
Cellular senescence - Mus musculus (house mouse)
PI3K-Akt signaling pathway - Mus musculus (house mouse)
p53 signaling pathway - Mus musculus (house mouse)
Small cell lung cancer - Mus musculus (house mouse)
Toxoplasmosis - Mus musculus (house mouse)
Yersinia infection - Mus musculus (house mouse)
Measles - Mus musculus (house mouse)
Hepatitis B - Mus musculus (house mouse)
Arginine biosynthesis - Mus musculus (house mouse)
Mismatch repair - Mus musculus (house mouse)
0.05
0.10
0.15
0.20
GeneRatio
pvalue
0.01
0.02
0.03
0.04
Count
1
2
3
4
Taste transduction - Mus musculus (house mouse)
Phospholipase D signaling pathway - Mus musculus (house mouse)
Non-homologous end-joining - Mus musculus (house mouse)
Thiamine metabolism - Mus musculus (house mouse)
Glycosaminoglycan biosynthesis - chondroitin sulfate / dermatan sulfate - Mus musculus (house mouse)
Glycosylphosphatidylinositol (GPI)-anchor biosynthesis - Mus musculus (house mouse)
SNARE interactions in vesicular transport - Mus musculus (house mouse)
DNA replication - Mus musculus (house mouse)
Aldosterone-regulated sodium reabsorption - Mus musculus (house mouse)
Various types of N-glycan biosynthesis - Mus musculus (house mouse)
0.05
0.06
0.07
0.08
0.09
0.10
GeneRatio
pvalue
0.02
0.04
0.06
0.08
Count
1.00
1.25
1.50
1.75
2.00

Figure S5. Gene expression profiles of DMR-mapped genes in response to LiTFSI and PFOA treatments. The data are represented as mean ± SEM of three independent replicates. ‘*’ for p-value < 0.05.

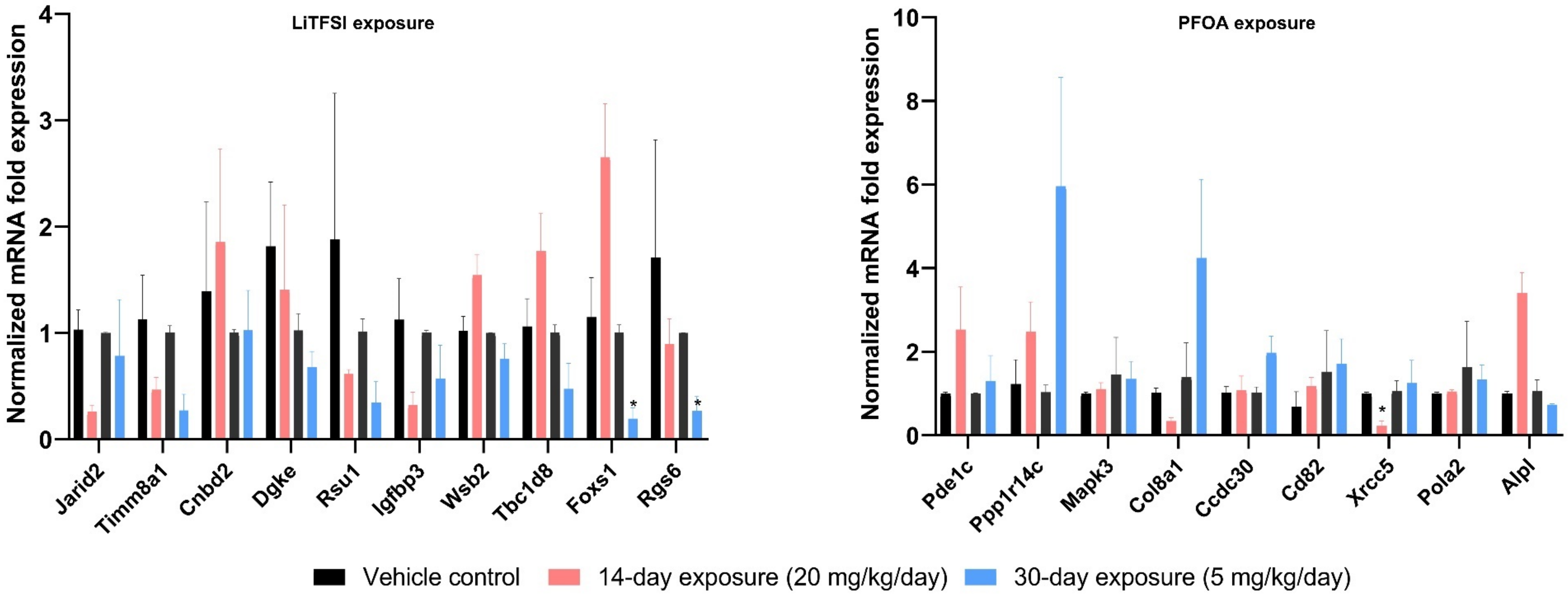

Table S1. Descriptive statistics of twelve methylomes before and after trimming.

| **Treatment** | **Sample** | **Number of Reads** | **Total Reads after trimming** | **Aligned Reads** | **Unaligned Reads** | **Ambiguously Aligned Reads** | **Total Cs** | **Methylated CpGs** | **Unmethylated CpGs** | **Methylated chgs** | **Unmethylated chgs** | **Methylated CHHs** | **Unmethylated CHHs** |
|---|---|---|---|---|---|---|---|---|---|---|---|---|---|
| **14-day_Control_1** | A2_kidney | 25,242,768 | 25096759 | 17843890 | 1759861 | 5493008 | 3.22E+08 | 10352701 | 47062270 | 383852 | 81312111 | 820142 | 1.82E+08 |
| **14-day_Control_2** | A3__kidney | 23,703,031 | 23556183 | 16023652 | 1874884 | 5657647 | 2.77E+08 | 9962902 | 38746090 | 341747 | 70214851 | 738319 | 1.57E+08 |
| **14-day_Control_3** | A4__kidney | 26,268,256 | 26123567 | 17772786 | 2194492 | 6156289 | 3.12E+08 | 10523585 | 43384442 | 362768 | 78905960 | 788492 | 1.78E+08 |
| **14-day_PFOA_20_1** | C2__kidney | 25,771,102 | 25615885 | 17980567 | 1775789 | 5859529 | 3.19E+08 | 10857548 | 44442931 | 389994 | 80643539 | 860859 | 1.81E+08 |
| **14-day_PFOA_20_2** | C3__kidney | 23,041,150 | 22900806 | 15462775 | 2253534 | 5184497 | 2.79E+08 | 10089267 | 36078864 | 326124 | 69669341 | 742433 | 1.62E+08 |
| **14-day_PFOA_20_3** | C4__kidney | 24,014,575 | 23799308 | 16315323 | 2199126 | 5284859 | 2.91E+08 | 8947092 | 41428372 | 288855 | 73098840 | 640545 | 1.67E+08 |
| **14-day_LITFSI_20_1** | E1__kidney | 28,710,269 | 28563753 | 19856864 | 2260924 | 6445965 | 3.55E+08 | 12467402 | 51041646 | 365814 | 89328188 | 806100 | 2.01E+08 |
| **14-day_LITFSI_20_2** | E2__kidney | 22,529,731 | 22406782 | 15250846 | 1867602 | 5288334 | 2.6E+08 | 8670294 | 38323445 | 270425 | 66378028 | 584878 | 1.46E+08 |
| **14-day_LITFSI_20_3** | E5__kidney | 25,712,198 | 25551282 | 17437609 | 1971128 | 6142545 | 3.09E+08 | 10548740 | 40920581 | 324503 | 78332293 | 708736 | 1.78E+08 |
| **30-day_Control_1** | F1__kidney | 25,719,499 | 25569890 | 17412783 | 2119710 | 6037397 | 3.09E+08 | 10392720 | 41502806 | 363428 | 77996017 | 797939 | 1.78E+08 |
| **30-day_Control_2** | F2__kidney | 23,704,872 | 23580868 | 16042597 | 1965822 | 5572449 | 2.72E+08 | 9180265 | 40037625 | 330163 | 69109707 | 693942 | 1.53E+08 |
| **30-day_Control_3** | F4__kidney | 26,776,048 | 26581827 | 17805860 | 2830422 | 5945545 | 3.17E+08 | 11714107 | 41863720 | 430172 | 78882273 | 950101 | 1.83E+08 |

| | | | | | | | | | | | | | |
|---|---|---|---|---|---|---|---|---|---|---|---|---|---|
| **30-day_PFOA_5_1** | H2__kidney | 24,554,994 | 24418020 | 16507589 | 2283257 | 5627174 | 2.83E+08 | 9627055 | 41536391 | 347913 | 71235001 | 765974 | 1.6E+08 |
| **30-day_PFOA_5_2** | H3__kidney | 24,945,594 | 24788633 | 16903097 | 2226182 | 5659354 | 3.08E+08 | 10549812 | 39412370 | 401660 | 77272467 | 903913 | 1.8E+08 |
| **30-day_PFOA_5_3** | H5__kidney | 25,668,212 | 25509071 | 17332007 | 2387268 | 5789796 | 3.14E+08 | 11320311 | 38672503 | 375528 | 78710306 | 851669 | 1.85E+08 |
| **30-day_LITFSI_5_1** | J1__kidney | 27,049,916 | 26877987 | 18183611 | 2414148 | 6280228 | 3.25E+08 | 12025028 | 41267437 | 385479 | 81462443 | 865066 | 1.89E+08 |
| **30-day_LITFSI_5_2** | J2__kidney | 26,560,439 | 26369958 | 17958155 | 2378645 | 6033158 | 3.28E+08 | 11775696 | 40446865 | 409590 | 81871915 | 924230 | 1.92E+08 |
| **30-day_LITFSI_5_3** | J5__kidney | 25,196,664 | 25038722 | 16850096 | 2592496 | 5596130 | 3.06E+08 | 11802788 | 38573496 | 406499 | 76545116 | 905192 | 1.78E+08 |

Table S2. Descriptive statistics of twelve methylomes before and after trimming

| Treatment | Sample Name | % GC | % Aligned | % Dups | Total Sequences(millions) | Number of CpG site |
|---|---|---|---|---|---|---|
| **14-day_Control_1** | A2_kidney | 34 | 71.1 | 82.2 | 25.1 | 1445553 |
| **14-day_Control_2** | A3__kidney | 34 | 68.4 | 82.9 | 23.6 | 1303050 |
| **14-day_Control_3** | A4__kidney | 33 | 68.9 | 83.2 | 26.1 | 1364531 |
| **14-day_PFOA_20_1** | C2__kidney | 34 | 71.3 | 82.6 | 25.6 | 1364531 |
| **14-day_PFOA_20_2** | C3__kidney | 33 | 69.0 | 81.9 | 22.9 | 1399176 |
| **14-day_PFOA_20_3** | C4__kidney | 33 | 67.9 | 82.4 | 23.8 | 1409566 |
| **14-day_LITFSI_20_1** | E1__kidney | 34 | 68.8 | 84.0 | 28.6 | 1454225 |
| **14-day_LITFSI_20_2** | E2__kidney | 34 | 72.4 | 82.7 | 22.4 | 1262612 |
| **14-day_LITFSI_20_3** | E5__kidney | 33 | 72.5 | 82.8 | 25.6 | 1410181 |
| **30-day_Control_1** | F1__kidney | 33 | 68.0 | 82.7 | 25.6 | 1417644 |
| **30-day_Control_2** | F2__kidney | 34 | 71.3 | 82.9 | 23.6 | 1263183 |
| **30-day_Control_3** | F4__kidney | 33 | 68.6 | 82.6 | 26.6 | 1479445 |
| **30-day_PFOA_5_1** | H2__kidney | 34 | 63.1 | 82.8 | 24.4 | 1319231 |
| **30-day_PFOA_5_2** | H3__kidney | 33 | 66.0 | 81.8 | 24.8 | 1483073 |
| **30-day_PFOA_5_3** | H5__kidney | 33 | 65.0 | 81.9 | 25.5 | 1497634 |
| **30-day_LITFSI_5_1** | J1__kidney | 33 | 64.7 | 83.0 | 26.9 | 1460767 |
| **30-day_LITFSI_5_2** | J2__kidney | 33 | 73.0 | 82.3 | 26.4 | 1506921 |
| **30-day_LITFSI_5_3** | J5__kidney | 33 | 68.0 | 81.8 | 25 | 1469873 |

Table S3. Top 10 genes mapped to DMRs of LiTFSI and PFOA -exposed kidneys. **The results did not include DMRs in intergenic regions.*

| Chr | Region | Gene Symbol | Gene Name | Methylation change | q value | CGIs |
|---|---|---|---|---|---|---|
| **LiTFSI-14-days-male** | | | | | | |
| Chr13 | Intron | Jarid2 | Jumonji, AT rich interactive domain 2 | 32.45 | 9.13E-105 | inter |
| Chr8 | promoter exon | Slc25a4 | Solute carrier family 25 | -26.76 | 2.65E-89 | inter |
| ChrX | promoter exon | Timm8a1 | translocase of inner mitochondrial membrane 8A1 | -36.50 | 8.18E-60 | inter |
| Chr6 | exon | Pde1c | Phosphodiesterase 1C | 25.88 | 7.48E-55 | inter |
| Chr2 | intron | Cnbd2 | Cyclic nucleotide binding domain containing 2 | 31.51 | 6.34E-51 | inter |
| Chr11 | intron | Igfbp3 | Insulin-like growth factor binding protein 3 | -59.32 | 5.50E-44 | inter |
| Chr5 | exon intron | Wsb2 | WD repeat and SOCS box-containing 2 | 61.87 | 4.19E-38 | inter |
| Chr1 | intron | Tbc1d8 | TBC1 domain family, member 8 | 64.18 | 5.82E-33 | inter |
| Chr11 | exon | Dgkeos | Diacylglycerol kinase, epsilon | -50.30 | 1.7E-27 | inter |
| Chr2 | exon | Foxs1 | Forkhead box S1 | 38.12 | 1.07E-24 | inter |
| **LiTFSI-30-day-male** | | | | | | |
| chr13 | intron | Jarid2 | Jumonji, AT rich interactive domain 2 | -28.41 | 5.6E-108 | inter |
| chrX | promoter exon | Timm8a1 | translocase of inner mitochondrial membrane 8A1 | -33.75 | 3.03E-74 | inter |
| chr14 | intron | Nrg3 | Neuregulin 3 | 29.78 | 5.4E-61 | inter |
| chr13 | intron | Jarid2 | Jumonji, AT rich interactive domain 2 | -48.18 | 1.47E-56 | inter |
| chr19 | exon | Vps13a | Vacuolar protein sorting 13A | -50.34 | 8.4E-49 | inter |
| chr1 | exon | Rcsd1 | RCSD domain containing 1 | -29.48 | 4.08E-43 | inter |
| chr12 | intron | Rgs6 | Regulator of G-protein signaling 6 | -49.26 | 3.26E-42 | inter |
| chr2 | intron | Cnbd2 | Cyclic nucleotide binding domain containing 2 | -29.69 | 4.46E-42 | inter |
| chr12 | exon | D430020J02Rik | RIKEN cDNA D430020J02 gene | 44.95 | 3.37E-41 | inter |
| chr4 | intron | Alpl | lkaline phosphatase, liver/bone/kidney | 51.61 | 2.76E-37 | inter |
| chr13 | promoter | Rpl17-ps6 | ribosomal protein L17, pseudogene 6 | 56.40 | 3.65E-34 | inter |
| **PFOA-14-day-male** | | | | | | |
| chr6 | exon | Pde1c | Phosphodiesterase 1C | 28.62 | 2.43E-88 | inter |
| chr6 | exon | Pde1c | Phosphodiesterase 1C | 28.09 | 9.12E-87 | inter |
| chr10 | promoter exon | Ppp1r14c | Protein phosphatase 1, regulatory inhibitor subunit 14C | -26.18 | 1.58E-77 | inter |
| chr7 | promoter exon | Mapk3 | Mitogen-activated protein kinase 3 | 26.34 | 6.83E-70 | inter |

| | | | | | | |
|---|---|---|---|---|---|---|
| chrX | promoter exon | Timm8a1 | translocase of inner mitochondrial membrane 8A1 | -36.34 | 6.49E-69 | inter |
| chr5 | exon intron | Wsb2 | WD repeat and SOCS box-containing 2 | 62.87 | 6.27E-55 | inter |
| chr2 | intron | Pcsk2 | Proprotein convertase subtilisin/kexin type 2 | 76.22 | 1.02E-52 | inter |
| chr16 | exon | Col8a1 | collagen, type VIII, alpha 1 | -63.81 | 4.29E-43 | inter |
| chr9 | intron | Rexo2 | RNA exonuclease 2 | 57.18 | 3.96E-42 | inter |
| chr13 | intron | Jarid2 | Jumonji, AT rich interactive domain 2 | 42.22 | 5.30E-35 | inter |
| chr4 | exon | Ccdc30 | coiled-coil domain containing 30 | 32.69 | 1.15E-29 | inter |
| **PFOA-10-day -female** | | | | | | |
| chr6 | intron | Tbxas1 | Thromboxane A synthase 1, platelet | 43.61 | 6.15E-120 | inter |
| chr13 | intron | Zfp131 | Zinc finger protein 131 | -25.99 | 7.91E-118 | inter |
| chr17 | intron | Wiz | Widely interspaced zinc finger motifs | -62.61 | 3.17E-113 | Shelves |
| chr4 | intron | St3gal3 | ST3 beta-galactoside alpha-2,3-sialyltransferase 3 | 70.52 | 2.33E-112 | inter |
| chr12 | intron | Numb | NUMB endocytic adaptor protein | 70.86 | 2.85E-111 | inter |
| chr17 | intron | Notch4 | Notch 4 | -68.53 | 2.19E-101 | inter |
| chr5 | intron | Slc15a4 | Solute carrier family 15, member 4 | -35.08 | 1.39E-94 | inter |
| chr7 | intron | Katnip | Katanin interacting protein | -46.13 | 3.15E-94 | inter |
| chr3 | intron | Slc44a5 | Solute carrier family 44, member 5 | -50.52 | 1.40E-90 | inter |
| chr13 | intron | Arhgef28 | Rho guanine nucleotide exchange factor (GEF) 28 | -65.36 | 2.66E-88 | inter |
| **PFOA-30-day -male** | | | | | | |
| chr2 | intron | Cd82 | CD82 antigen | -79.34 | 1.00E-94 | inter |
| chr8 | intron | Kbtbd11 | kelch repeat and BTB (POZ) domain containing 11 | -77.33 | 5.31E-94 | inter |
| chr2 | intron | Ldlrad3 | low density lipoprotein receptor class A domain containing 3 | 63.57 | 7.40E-52 | inter |
| chr2 | intron | Alx4 | aristaless-like homeobox 4 | -38.09 | 3.96E-50 | inter |
| chr12 | exon | D430020J02Rik | RIKEN cDNA D430020J02 gene | 50.79 | 9.24E-44 | inter |
| chr1 | intron | Xrcc5 | X-ray repair complementing defective repair in Chinese hamster cells 5 | -59.27 | 2.00E-43 | inter |
| chr19 | exon | Pola2 | polymerase (DNA directed), alpha 2 | -43.36 | 3.87E-42 | shore |
| chr4 | intron | Alpl | alkaline phosphatase, liver/bone/kidney | 54.97 | 2.21E-40 | inter |

| | | | | | | |
|---|---|---|---|---|---|---|
| chr11 | promoter exon | Gm11620 | predicted gene 11620 | -31.53 | 5.02E-39 | inter |
| chr13 | promoter | Rpl17-ps6 | ribosomal protein L17, pseudogene 6 | 62.036 | 1.18E-38 | inter |

Table S4. Genes mapped to DMRs overlaps in 14-day and 30-day LiTFSI and PFOA -exposed kidney samples respectively.

| Chr | Region | Gene Symbol | Gene name | 14-day meth change | 30-day meth change | CGIs |
|---|---|---|---|---|---|---|
| **LiTFSI- -male** | | | | | | |
| Chr13:45023101-45023200 | intron | Jarid2 | Jumonji, AT rich interactive domain 2 | 32.5 | -28.41 | inter |
| chr13: 45023201-45023300 | intron | Jarid2 | Jumonji, AT rich interactive domain 2 | 34.9 | -48.18 | inter |
| ChrX:133442301-133442400 | promoter exon | Timm8a1 | translocase of inner mitochondrial membrane 8A1 | -36.5 | -33.75 | inter |
| chr2:156204901-156205000 | intron | Cnbd2 | Cyclic nucleotide binding domain containing 2 | 31.5 | -29.69 | inter |
| chr10: 11367801-11367900 | intergenic | Gm48679 | Predicted gene 48679 | -29.3 | -28.24 | inter |
| chr11: 89005501-89005600 | intergenic | Gm11497 | Predicted gene 11497 | -43.2 | 39.40 | inter |
| chr12: 116139501-116139600 | intergenic | Gm23732 | Predicted gene 23732 | 53.4 | 36.28 | inter |
| chr11: 88929901-88930000 | exon | Dgkeos | Diacylglycerol kinase, epsilon | -50.3 | 34.03 | inter |
| chr2: 13176701-13176800 | intron | Rsu1 | Ras suppressor protein 1 | -35.4 | 30.20 | inter |
| chr15: 100218301-100218400 | intron | Mettl7a3 | Methyltransferase like 7A3 | -37.3 | 28.20 | inter |
| chr11: 116179301-116179400 | exon | Exoc7 | Exocyst complex component 7 | 31.6 | -33.38 | inter |
| chr7: 136729701-136729800 | exon | Mgmt | O-6-methylguanine-DNA methyltransferase | 33.9 | -40.61 | inter |
| chr5: 76970701-76970800 | intron | Cracd | Cysteine-rich acidic secretory | -27.3 | 56.55 | inter |

| | | | | | | |
|---|---|---|---|---|---|---|
| | | | protein D | | | |
| chr9: 82993501-82993600 | intron | Hmgn3 | High-mobility group nucleosome-binding domain-containing protein 3 | -30.5 | 38.92 | inter |
| chr12: 113153801-113153900 | intergenic | Tmem121 | Transmembrane protein 121 | 28.9 | 33.58 | inter |
| chr8: 15066301-15066400 | intron | Kbtbd11 | Kelch repeat and BTB domain-containing protein 11 | 30.0 | -47.36 | inter |
| chr7: 31153701-31153800 | intergenic | Scgb1b28-ps | Secretoglobin family 1B member 28, pseudogene | 38.0 | -38.86 | inter |
| chr15: 88976201-88976300 | intron | Selenoo | Selenoprotein O | -29.4 | 42.52 | inter |
| chr3: 30432101-30432200 | intron | Mecom | MDS1 and EVI1 complex locus protein EVI1 | -26.6 | -44.18 | inter |
| chr5:4944601-4944700 | intron | Cdk14 | Cyclin-dependent kinase 14 | 32.1 | 27.96 | inter |
| chr2: 129843501-129843600 | intergenic | Tgm3 | Transglutaminase 3 | -30.3 | 28.25 | inter |
| chr1: 105067601-105067700 | intergenic | Gm29012 | Gene model 29012 | -33.1 | 25.32 | inter |
| chr5: 114934601-114934700 | intron | Ankrd13a | Ankyrin repeat domain-containing protein 13A | -27.5 | 26.54 | shelves |
| chr7: 110470501-110470600 | exon | Irag1 | Interferon regulatory factor-associated protein 1 | -29.4 | 36.04 | inter |
| **PFOA- -male** | | | | | | |
| chr15: 45185201-45185300 | intergenic | Gm53014 | Jumonji, AT rich interactive domain 2 | 43.97 | 28.33 | inter |

| | | | | | | |
|---|---|---|---|---|---|---|
| chr19: 6003801-6003900 | exon | Pola2 | translocase of inner mitochondrial membrane 8A1 | 27.55 | -43.36 | shores |
| chr6: 92706101-92706200 | intergenic | Prickle2 | Prickle planar cell polarity protein 2 | -52.60 | 32.84 | inter |
| chr1: 84280201-84280300 | intron | Pid1 | Phosphotyrosine interaction domain-containing protein 1 | -44.69 | 46.15 | inter |
| chr15: 44821701-44821800 | promoter | Gm18153 | Gene model 18153 | -39.85 | -29.94 | inter |
| chr4: 149565101-149565200 | intron | Tmem274 | Transmembrane protein 274 | -35.89 | 37.01 | inter |
| chr15: 3297401-3297500 | promoter | Selenop | Selenoprotein P | 39.29 | 32.31 | inter |
| chr1: 90130101-90130200 | intergenic | Ackr3 | Atypical chemokine receptor 3 | -27.90 | 27.74 | inter |
| chr5: 144472101-144472200 | intergenic | Nptx2 | Neuronal pentraxin-2 | 28.14 | -65.99 | inter |
| chr11: 6797401-6797500 | intergenic | Gm11983 | Gene model 11983 | -39.84 | -27.21 | inter |
| chr3: 119892701-119892800 | intergenic | Gm18384 | Gene model 18384 | 32.92 | -38.47 | inter |
| chr1: 105067601-105067700 | intergenic | Gm29012 | Gene model 29012 | 28.05 | 31.73 | inter |
| chr5: 148731501-148731600 | intergenic | 4930505K14Rik | RIKEN cDNA 4930505K14 gene | -30.65 | 48.02 | inter |
| chrX: 89212501-89212600 | intergenic | Gm6027 | Gene model 6027 | -33.71 | -30.64 | inter |
| chr5: 76970701-76970800 | intron | Cracd | Cysteine-rich acidic secretory protein D | -25.86 | 41.76 | inter |
| chr6: 68347001-68347100 | intergenic | Gm5310 | Gene model 5310 | -29.67 | 32.94 | inter |
| chr5: 73500401-73500500 | intergenic | Ociad2 | OCIA domain-containing protein 2 | -27.15 | -35.90 | inter |

| | | | | | | |
|---|---|---|---|---|---|---|
| chr15: 95612901-95613000 | promoter | Gm8843 | Gene model 8843 | 27.24 | 27.19 | inter |
| chr7: 110317701-110317800 | intergenic | Gm18907 | Gene model 18907 | 32.21 | -58.16 | inter |
| chr19: 5992201-5992300 | intron | Pola2 | DNA polymerase alpha subunit B | 26.46 | -31.79 | shelves |
| chr11: 7159301-7159400 | intron | Igfbp3 | Insulin-like growth factor-binding protein 3 | -25.43 | -40.51 | inter |
| chr14: 34416001-34416100 | intron | Wapl | Wings apart-like protein homolog | -27.95 | 30.53 | shelves |
| chr10: 25427101-25427200 | intron | Gm29571 | Gene model 29571 | 28.56 | 51.97 | inter |
| chr15: 53739201-53739300 | intron | Gm7489 | Gene model 7489 | 29.73 | 44.14 | inter |
| chr7: 110470501-110470600 | exon | Irag1 | Interleukin-1 receptor-associated kinase 1 | -27.24 | 56.89 | inter |

Table S5. Genes mapped to DMRs overlaps in 14-day and 10-day PFOA -exposed kidney in male and female. **The results did not include DMRs in intergenic regions.*

| Chr | Region | Gene Symbol | Gene name | Male meth change | Female meth change | CGI |
|---|---|---|---|---|---|---|
| chr1: 30873301-30873400 | exon intron | Gm37522/ Phf3 | Jumonji, AT rich interactive domain 2 | -34.31 | -48.40 | inter |
| chr1: 118981301-118981400 | promoter exon | Gli2 | translocase of inner mitochondrial membrane 8A1 | 28.15 | 40.00 | inter |
| chr2: 18056501-18056600 | intergenic | Skida1 | Ski/Dach domain-containing protein 1 | -28.66 | -26.64 | shelves |
| chr2: 153241301-153241400 | exon | Gm14472 | Gene model 14472 | -28.94 | -26.15 | inter |
| chr2: 156863001-156863100 | exon | Soga1 | Sarcosine dehydrogenase [NAD(P)+], mitochondrial | -31.28 | -61.00 | inter |
| chr2: 156863201-156863300 | exon | Soga1 | Sarcosine dehydrogenase [NAD(P)+], mitochondrial | -25.67 | -64.67 | inter |
| chr3: 137867601-137867700 | intergenic | Mttp | Microsomal triglyceride transfer protein | -31.98 | -28.69 | inter |
| chr4: 117133601-117133700 | intron | Rnf220 | RING finger protein 220 | -29.92 | -39.97 | inter |
| chr4: 134868101-134868200 | intron | Runx3 | Runt-related transcription factor 3 | 29.64 | 42.59 | inter |
| chr4: 139380701-139380800 | promoter intron | Tas1r2/ Gm9870 | Taste receptor type 1 member 2 | -30.53 | -55.54 | inter |
| chr5: 73071301-73071400 | promoter exon | Slain2 | SLAIN motif-containing protein 2 | 28.21 | 29.03 | inter |
| chr5: 110378901-110379000 | promoter exon | Ankle2 | Ankyrin repeat and LEM domain-containing protein 2 | 31.59 | 51.69 | inter |

| | | | | | | |
|---|---|---|---|---|---|---|
| chr6: 48024601-48024700 | promoter exon | Gm54714/ Zfp777 | Gene model 54714 | 31.28 | 39.34 | inter |
| chr6: 83166001-83166100 | exon | Dctn1 | Dynactin subunit 1 | -28.77 | -42.74 | inter |
| chr6: 122819701-122819800 | exon intron | Foxj2 | Forkhead box protein J2 | -39.07 | -47.36 | inter |
| chr7: 24972001-24972100 | exon | Cic | Capicua transcriptional repressor | -27.96 | -41.17 | inter |
| chr8: 3587701-3587800 | exon intron | Pnpla6 | Patatin-like phospholipase domain-containing protein 6 | -28.16 | -36.44 | inter |
| chr8: 11547701-11547800 | promoter exon | Naxd | NAD(P)H-hydrate epimerase | 31.26 | 61.76 | shelves |
| chr8: 15012501-15012600 | promoter exon intron | Gm16350 | Gene model 16350 | -32.66 | -36.90 | inter |
| chr8: 83998001-83998100 | exon | Tbc1d9 | TBC1 domain family member 9 | -26.87 | -51.92 | inter |
| chr8: 125160701-125160800 | exon | Pgbd5 | Phosphoglycerate dehydrogenase pseudogene 5 | 28.79 | 47.61 | inter |
| chr9: 21829401-21829500 | promoter exon | Rab3d | Ras-related protein Rab-3D | 27.58 | 26.73 | inter |
| chr9: 121995901-121996000 | exon | Snrk | SNF-related serine/threonine-protein kinase | -25.85 | -50.34 | inter |
| chr10: 120869801-120869900 | promoter exon | Wif1 | WNT inhibitory factor 1 | 36.99 | 34.07 | inter |
| chr10: 126895301-126895400 | promoter exon | Marchf9 | E3 ubiquitin-protein ligase MARCH9 | 25.10 | 29.23 | inter |
| chr11: 62731701-62731800 | exon | Trim16 | Tripartite motif-containing protein 16 | -29.67 | -77.29 | inter |
| chr11: 98327601-98327700 | exon | Erbb2 | Receptor tyrosine-protein kinase erbB-2 | -29.97 | -39.21 | inter |

| | | | | | | |
|---|---|---|---|---|---|---|
| chr12: 108327801-108327900 | exon | Cyp46a1 | Cholesterol 24-hydroxylase | -28.09 | -52.70 | inter |
| chr13: 38151401-38151500 | exon intron | Ssr1 | Signal recognition particle subunit SRP54 | -27.29 | -60.64 | inter |
| chr13: 58401901-58402000 | intron | Gkap1 | Glycine-rich protein 1-associated kinase | -34.77 | -39.51 | inter |
| chr13: 110531501-110531600 | promoter exon | Plk2 | Serine/threonine-protein kinase PLK2 | 25.75 | 45.24 | inter |
| chr15: 57892301-57892400 | exon | 9130401M01Rik | RIKEN cDNA 9130401M01 gene | -25.82 | -33.91 | inter |
| chr15: 58823401-58823500 | exon | Ndufb9 | NADH dehydrogenase [ubiquinone] 1 beta subcomplex subunit 9 | -33.35 | -45.16 | inter |

Table S6. qPCR primer sequences used in this study.

| **Name** | **Sequence** |
|---|---|
| B2m-F | GCTACTCGGCGCTTCAGTCG |
| B2m-R | TCCCATTCTCCGGTGGGTGG |
| Tbp-F | GTCAGGCGTTCGGTGGATCG |
| Tbp-R | GCTGGGCACTGCGGAGAAAA |
| Jarid2-F | GGCTGCAGATGCTGGAGAGC |
| Jarid2-R | TCCCGGGAAGCAGACGACAA |
| Timm8a1-F | AGTTCCGGTTCCGTCACCCA |
| Timm8a1-R | AGCCCGAAGAGGACGTGGAG |
| Cnbd2-F | CTGGTGGCCTTGGGGAGGAT |
| Cnbd2-R | CCAGTTGCTGCCAGACCCAC |
| Dgke-F | TTGTGGAGGGGACGGGACTG |
| Dgke-R | AGCCTGTACCCCAACCCAGG |
| Rsu1-F | CACGCTGCCTCGAGGATTCG |
| Rsu1-R | TAGAGTGCACGCAGGGTGGT |
| Igfbp3-F | GAGTCCCAGAGGCGTCCACA |
| Igfbp3-R | AGCCCCGCTTTCTGCCTTTG |
| Wsb2-F | AGACGGTTCCTGGTTCGCCT |
| Wsb2-R | CCTTCAGACTGCCCCGTCCT |
| Tbc1d8-F | CTGGCAGATGTCACGCTCCG |
| Tbc1d8-R | TCTGTGCTCTGGCCTCCAGG |
| Foxs1-F | CAGCTACCAGTGCCGGATGC |
| Foxs1-R | CCCGGTGAGAGGCTGAAGGT |
| Rgs6-F | GGCGGGACCAGTTCCTCAGA |
| Rgs6-R | TCCTCCACCCTCTTGGCCAC |
| IL-6-F | TCTGGAGCCCACCAAGAACGA |
| IL-6-R | GGAAGGCCGTGGTTGTCACC |
| TNF-F | GGCCCAGACCCTCACACTCA |

| | |
|---|---|
| TNF-R | CCACTCCAGCTGCTCCTCCA |
| Tgfb1-F | ATCCCACCTTTGCCGAGGGT |
| Tgfb1-F | CGGGCGTCAGCACTAGAAGC |
| Ifng-F | TCCGAGTGGTCCACCAGCTG |
| Ifng-R | TTCCCCACCCCGAATCAGCA |
| ppara-F | GCCACTTCGAGTCCCCTGGA |
| ppara-R | CCTCCAGCCCCCAAAACAGC |
| Abcg2-F | CCATGGGCCAGCACAGAAGG |
| Abcg2-R | CCGCAGGGTTGTTGTAGGGC |
| Alpl-F | GCCTTCATAAGCAGGCGGGG |
| Alpl-R | CGATCCTGCAGGCACTCGTG |
| Ccdc30-F | CCAAGTCCCCCGAGCCTCTT |
| Ccdc30-R | TGCTCCTGCTCGGTTTGCAC |
| Cd82-F | CGCGGGACTTAAAGCGCGTA |
| Cd82-R | CTTTGACACAGCCTGCCCCC |
| Col8a1-F | GGGCTGCTTGGACCCAAAGG |
| Col8a1-R | CGGAATCCCAGGGGGTCCAA |
| Mapk3-F | TCCCACTCCAATCCCCTGCC |
| Mapk3-R | CACCCCTCCGTAGCACAGGT |
| Pde1c-F | TTGCTCGCTTCTCCCGGTCT |
| Pde1c-R | TGGGCTATCCAGCTGTGGCA |
| Pola2-F | TCCACCCCACGAGGACATGG |
| Pola2-R | TGCCGCCCACTTGTCCTTTG |
| Ppp1r14c-F | TCGCGTCTGCAGGGCAAAAA |
| Ppp1r14c-R | TCCAAGGACCCTGGTGCTCC |
| Slc2a9-F | CAGGACCAAGCTCCCTGGGT |
| Slc2a9-R | TGCGATCATGAAGGCTGCCG |
| Slc22a6-F | TGTGTGCTCTCATCGGGCCT |
| Slc22a6-R | GATCAGCACCTTCCGACGGC |
| Slc22a8-F | accctgcctcaggtcactcg |

| | |
|---|---|
| Slc22a8-R | GCCCATGCTTCCAACACGGT |
| Slc22a12-F | CCTGAGACCCATGGCCCAGT |
| Slc22a12-R | AGAGGGGGAAGGTGGGCATG |
